# Development of DARPin T cell engagers for specific targeting of tumor-associated HLA/peptide complexes

**DOI:** 10.1101/2025.03.28.645643

**Authors:** Natalia Venetz-Arenas, Tim Schulte, Sandra Müller, Karin Wallden, Stefanie Fischer, Tom Resink, Nadir Kadri, Maria Paladino, Nicole Pina, Filip Radom, Denis Villemagne, Sandra Bruckmaier, Andreas Cornelius, Tanja Hospodarsch, Evren Alici, Hans-Gustaf Ljunggren, Benedict J. Chambers, Xiao Han, Renhua Sun, Marta Carroni, Victor Levitsky, Tatyana Sandalova, Marcel Walser, Adnane Achour

## Abstract

The compromise between affinity and specificity in TCR-dependent targeting of HLA-restricted tumor-associated antigens presents a significant challenge in developing efficacious immunotherapies. As such, T cell engagers which circumvent these limitations are of particular interest. We have established a process to generate bispecific Designed Ankyrin Repeat Proteins (DARPins) that simultaneously target HLA-I molecules in complex with tumor-associated peptides and CD3ε. High-affinity HLA-A*0201/NY-ESO1_157-165_-specific DARPins were isolated after only four rounds of *in-vitro* selection from naïve DARPin libraries. Combining HLA-A*0201/NY-ESO1_157-165_-specific DARPins with a CD3ε-specific DARPin created potent T cell engagers which elicited CD8^+^ T cell activation towards tumor targets with high peptide specificity, as confirmed by X-scanning mutagenesis and functional killing assays. The cryo-EM structure of a ternary DARPin/HLA-A*0201/NY-ESO1_157-165_ complex revealed a rigid and concave DARPin surface that binds to the entire length of the peptide-binding cleft, contacting both α-helices and the peptide. The present results unveil promising immuno-oncotherapeutic approaches with the possibility of rapidly developing DARPins with high affinity and specificity to HLA/peptide targets that can be readily combined with a new generation of anti-CD3ε-specific DARPins.

## Introduction

Technological advances in mass-spectrometry and other technologies have significantly enhanced the capacity to identify and map HLA-restricted tumour-associated antigens (TAA) and neoantigen repertoires that can be used as targets for CD8^+^ cytotoxic T lymphocyte (CTL)-mediated cancer immunotherapies. The further possibility to direct adaptive immune responses through strategies, such as peptide vaccination ^1–5^ or adoptive T cell transfer,^6,7^ has provided promising results in several malignancies. ^11–14^ In particular, the design of TCR-like antibodies, modified TCRs, or various T cell engagers (TCEs) with enhanced affinities to tumor-specific HLA/peptide complexes allows for individualized therapeutic approaches.^7–9^ For example, engineered high-affinity, soluble TCRs specific to the melanoma-associated antigen gp100 or the more widely cancer-associated NY-ESO1 peptides have shown success in preclinical models and clinical trials with regard to specificity and potency.^10–14^ Even with these clinical successes, many patients do not develop adequate responses, acquire resistance, or both.^15^ Therefore, an unexplored potential to further enhance clinical response rates for patients through additional novel approaches remains.

Despite considerable progress over the last 30 years, the exact molecular bases governing specific activation of CD8^+^ T cell responses remain partially solved, impeding the widespread development of peptide-based immunotherapies. Moreover, T cell tolerance must be considered as many TAAs are derived from endogenous proteins. The lack of therapeutic efficacy in some previous studies could be attributed to the induction of tolerance in the available T cell repertoire towards HLA-I in complex with dominant TAAs.^16–18^ Additionally, TCRs are relatively weak binders that recognize HLA alleles in complex with a limited pool of peptides, when compared with high-affinity, antigen-specific antibodies.^8,19^ Notwithstanding the affinity-limiting effect of thymic deselection on displayed TCRs of HLA/TAA restricted T cell populations, these populations may be further functionally suppressed by the tumor microenvironment or exhausted due to cancer immunoediting.^20,21^ In practice, the weaker affinity of TCRs can be overcome through *in vitro* selection and affinity maturation strategies to obtain affinity-matured TCRs, TCR-like antibody binders,^7,22^ or altered peptide ligands.^23–27^ Although TCRs can display exceptional specificity to an HLA/peptide complex with high sensitivity to conservative mutations, the same TCRs may also bind multiple other HLA/peptide complexes through cross-recognition, potentially leading to unwanted outcomes.^28,29^ This nonspecific recognition can be exacerbated by gains in affinity,^30,31^ as exemplified by the high-affinity TCR developed against the cancer-associated HLA-A*0101/MAGE-A3 target. Clinical administration of T cells expressing the affinity-enhanced TCR, which displayed promising *in vitro* specificity, induced lethal off-target responses against an HLA-A*0101-restricted, titin-derived peptide on cardiomyocytes.^32,33^

We reasoned that designed ankyrin repeat proteins (DARPins) might be particularly suited as alternative antigen recognition molecules to overcome the aforementioned limitations. The intrinsic rigidity of DARPins, combined with their longitudinal and transversal size,^34,35^ led us to hypothesize that these binding molecules could effectively engage the entire length of the peptide binding cleft of an HLA/peptide target molecule. The versatility of DARPins, combined with their high affinity and specificity, resulted into their successful use as therapeutics against a wide variety of targets.^36–38^ DARPins, which are based on a ankyrin repeat scaffold, create an elongated solenoid fold that forms a rigid concave interface to bind target molecules.^34,39,40^ In contrast to antibodies and TCRs, DARPins do not contain cysteine residues. Therefore, they can both be easily expressed in soluble form in *Escherichia coli* at very high yields and possess favorable biophysical properties.^41,42^

By combining two DARPin molecules with high specificity to HLA-A*0201/NY-ESO1_157-165_ and CD3ε, respectively, we developed bispecific DARPin TCE constructs that produce potent and specific CD8^+^ T cell activation in the presence of HLA-A*0201/NY-ESO-1_157-165_-positive tumor cells. Further affinity maturation of the CD3ε-binding DARPin allowed us to enhance the potency of the identified bispecific DARPin TCEs without compromising their specificity. The cryo-EM structure of the DARPin lead candidate in complex with HLA-A*0201/NY-ESO1_157-165_ revealed that the DARPin binding surface spans the entire length of the HLA peptide binding cleft contacting both helixes and peptide. As a result, DARPins, such as those developed here, are a highly attractive alternative to the more commonly employed engineered TCR or TCR-mimicking antibodies for specific and potent targeting of HLA/peptide complexes.

## Results

### Identification and validation of DARPins with high affinity and specificity to HLA-A*0201/NY-ESO1_157-165_

We initiated this study by selecting HLA-A*0201/NY-ESO1_157–165_(9V) (SLLMWITQ<u>V</u>) as a target, given that modification of the anchor residue p9C in HLA-A*0201-restricted NY-ESO1_157165_ (SLLMWITQ<u>C</u>) to valine significantly improves the HLA/peptide complex stability and TCR recognition without altering peptide conformation.^23^ Using four different DARPin ribosome display mRNA libraries with physical diversities of approximately 10^12^ variants, we performed four rounds of selection and counter-selection against HLA-A*0201/NY-ESO1_157-165_(9V) and the negative control HLA-A*0201/EBNA1_562-570_ (FMVFLQTHI), respectively (**Figure 1A**). Selected DARPin candidates were screened using homogenous time-resolved FRET (HTRF) to detect binding to HLA-A*0201/NY-ESO1_157–165_(9V) (**Figure 1B**). Off-target binding was assessed using the negative controls HLA-A*0201/EBNA1_562-570_ and HLA-A*0201/NY-ESO1_157-165_(9VAA) (SLL<u>AA</u>ITQV), in which the major TCR-interacting peptide residues p4M and p5W ^43^ were mutated to alanine. All DARPin binders that generated >50-times higher HTRF signals towards HLA-A*0201/NY-ESO1_157-165_(9V) compared to both negative controls were considered specific hits. Each identified hit was sequenced, cloned, and produced in a bispecific TCE format by linking the HLA-A*0201/NY-ESO1_157-165_(9V)-specific DARPin binder to a recently developed anti-CD3ε DARPin candidate ^44^ (**Figure 1C**, Supplementary Table 1).

**Figure 1.**
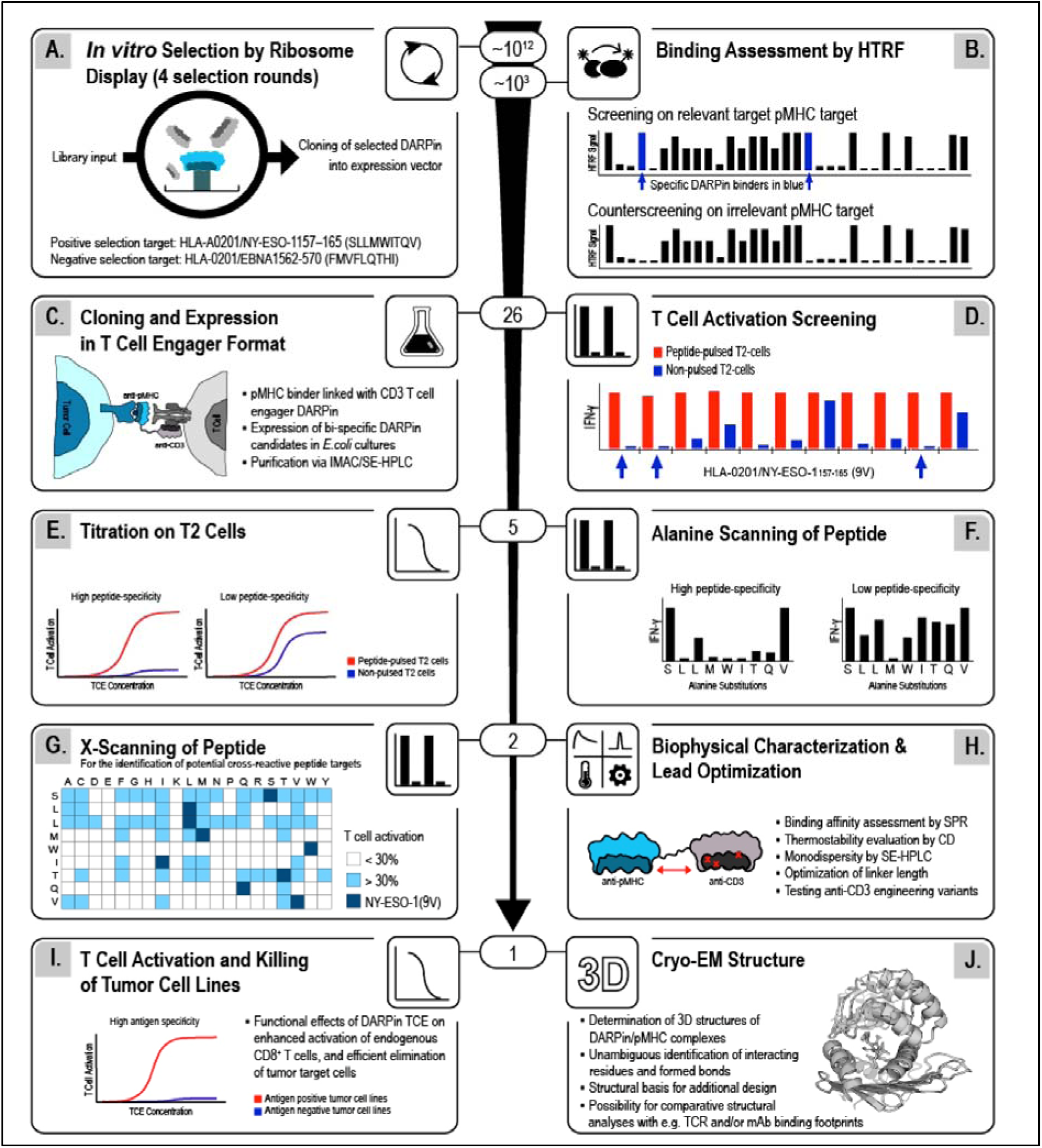
Generation of HLA/peptide-specific DARPins A general overview for the generation and selection of bispecific DARPin T cell engagers (TCEs) is presented with focus on HLA-A*0201/NY-ESO1_157-165_ as proof of concept. **A.** A physical DARPin library, with a diversity of approximately 10^12^ complexes, was used to perform *in vitro* ribosome display selection of HLA-A*0201/NY-ESO1_157-165_(9V)-specific DARPin binders. Negative selection was performed using HLA-A*0201/EBNA1_562-570_. **B.** Homogenous Time-Resolved FRET (HTRF) was used to screen the binding capacity of approximately 1000 selected DARPin candidates to HLA-A*0201/NY-ESO1_157-165_(9V). HLA-A*0201/EBNA1_562-570_ was used as negative control. **C.** A total of 26 HLA-A*0201/NY-ESO1_157-165_(9V)-specific DARPin binders were selected, and DARPin TCEs were created by linking each of them to a CD3ε-specific DARPin. **D.** T-cell activation screenings, performed with peptide-pulsed or non-pulsed T2 cells in the presence of CD8 T cells and three different concentrations of DARPin TCEs, allowed the identification of five DARPin TCE top candidates. See **Supplementary Figure 1**. **E.** The specificity and potency of DARPin TCEs were further verified using a panel of HLA-A*0201^+^ tumor cells that express the wild-type epitope NY-ESO1_157-165_(9C). See Figure 2. **F.** Alanine scanning mutation covering all peptide positions revealed the specificity of each DARPin TCE to the entire peptide space. See Figure 2C. **G.** X-scanning T-cell activation assay to identify possible substitution residues at each peptide position. The investigated peptide foot-print matrix was entered into the Expasy database for the identification of potential cross-reactive human peptide-HLA complexes. See Figure 3. **H.** Biophysical characterization of the two lead DARPin TCE candidates and evaluation of the effects of various linker lengths and CD3ε-specific DARPin variants. See Figure 4 **and Supplementary Figure 2**. **I.** The remaining candidate was screened in T-cell activation and killing assays against a panel of antigen-positive and antigen-negative tumor cell lines. See Figure 5. **J.** The three-dimensional structure of the final lead DARPin candidate in complex with HLA-A*0201/NY-ESO1_157–165_(9V) was determined to ∼3 Å resolution using cryo-electron microscopy. See Figure 6.

### Development and validation of bispecific DARPin TCEs targeting HLA-A*0201/NY-ESO1_157-165_ and CD3ε

The capacity of the 26 different DARPin TCEs for mediating highly specific and potent activation of CD8^+^ T cells, from peripheral blood mononuclear cells (PBMCs) from healthy donors, was screened. To this end, intracellular IFNγ levels were measured in the presence of either NY-ESO1_157-165_(9V)-pulsed TAP-deficient T2 cells or non-pulsed T2 cells (**Figure 1D, Supplementary Figure 1**). Subsequently, the ability of five select DARPin TCEs to elicit wild-type NY-ESO1_157-165_(9C)-specific activation of CD8^+^ T cells was assessed by co-incubating T cells and each TCE with HLA-A*0201^+^ tumor target cell lines expressing NY-ESO1_157-165_(9C) (**Figure 1E**). This demonstrated the unambiguous capacity of five selected DARPin TCEs to recognize the wild-type epitope in the presence of physiological target ligand levels. Notably, the five lead DARPin TCE candidates, henceforth referred to as NY_1xCD3 to NY_5xCD3, provoked a significant dose-dependent increase of the CD8^+^ T cell activation marker CD25 (**Figure 2A**) and IFNγ production by CD8^+^ T cells (**Figure 2B)** in the presence of HLA-A*0201^+^/NY-ESO1_157-165_(9C)-positive tumor cells. In contrast, CD25 and IFNγ expression levels remained insignificant following the co-incubation of CD8^+^ T cells with NY-ESO1-negative tumor target cells in the presence of most tested DARPin TCEs. The capacity of each DARPin TCE candidate to elicit CD8^+^ T cell responses was also assessed with T2 cells pulsed with single alanine variants of NY-ESO1_157-165_ (**Figures 1F and 2C)**. The results indicate that the DARPin TCE lead candidates NY_1xCD3, NY_2xCD3, and NY_3xCD3 display a high specificity for most peptide positions (**Figure 2C**). NY_4xCD3 and NY_5xCD3 DARPins, however, both provoked unspecific cytokine production (**Figures 2A and 2B**) and were only sensitive to mutations at peptide residues 4-6 and 5-6, respectively (**Figure 2C**). We decided to proceed further with NY_1xCD3 and NY_2xCD3 which displayed small but significant differences in their peptide specificities (**Figure 2C)**. NY_3xCD3 was discarded given the similarities in peptide activation profiles when compared with NY_2xCD3 (**Figure 2C**).

**Figure 2.**
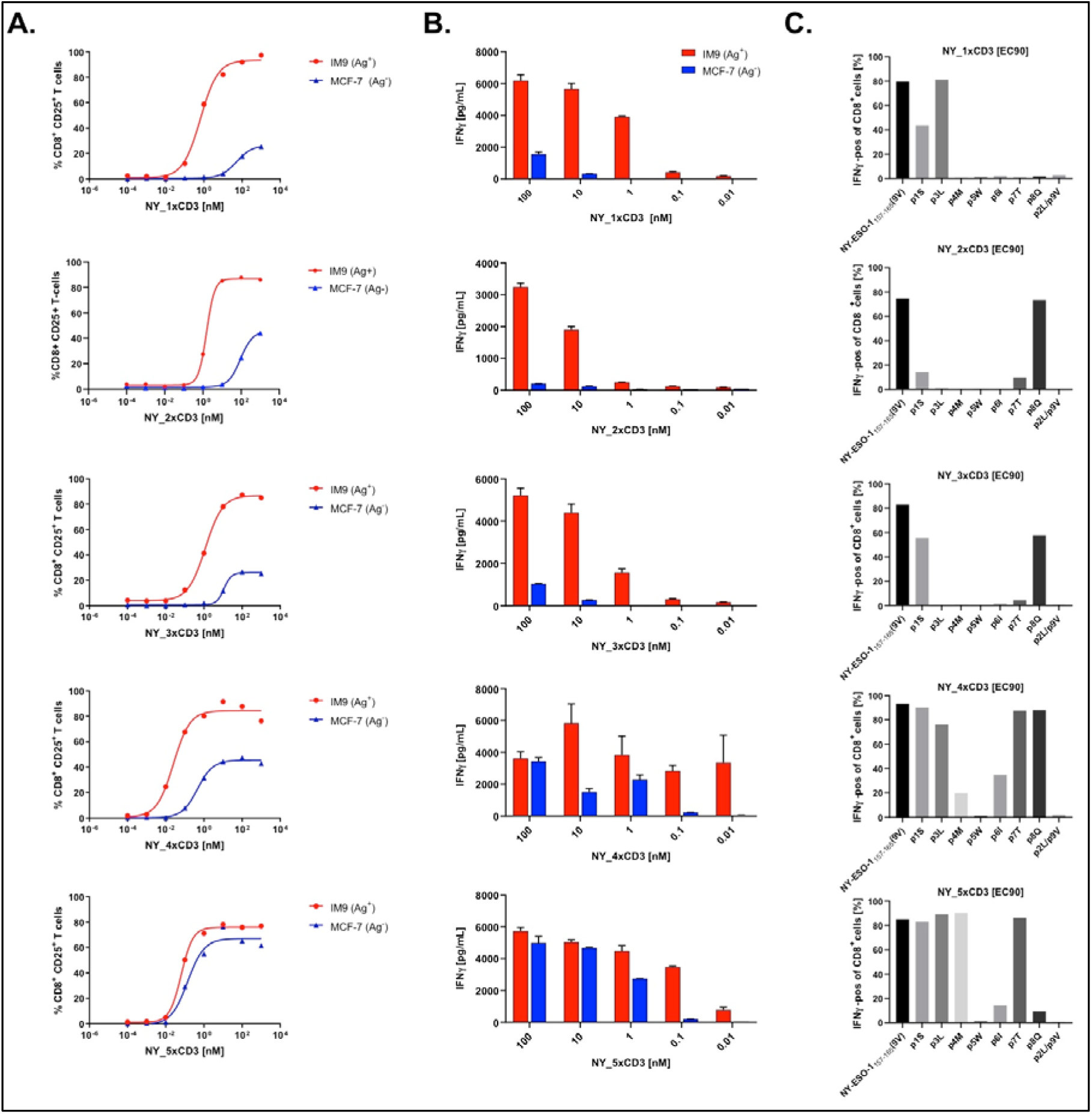
DARPin TCEs provoke significant CD8^+^ T cell responses with high specificity to HLA-A*0201/NY–ESO1_157-165_ **A-B.** The HLA-A*0201^+^/NY-ESO1^+^ (Ag^+^) and HLA-A*0201^+^/NY-ESO1^-^ (Ag^-^) tumor cell lines IM9 and MCF-7, respectively, were incubated with PBMCs for 48 hours in the presence or absence of each DARPin TCE. T cell activation was assessed by measuring the levels of the activation marker CD25 on CD8^+^ T cells (**A**) and IFNγ release (**B**). **C**. T2 cells, pulsed with 1μM alanine-mutated peptide variants of NY-ESO1_157–165_(9V), were incubated with effector CD8^+^ T cells and each DARPin TCE, and the amount of IFNγ positive CD8^+^ T cells was assessed. While NY_1xCD3, NY_2xCD3 and NY_3xCD3 were all highly sensitive to mutations throughout the entire peptide length, NY_4xCD3 and NY_5xCD3 were mainly sensitive to modifications only at peptide residues p4-p6. Results are representative of at least three independent experiments.

HPLC-size exclusion chromatography analyses confirmed both lead TCE candidates are monodisperse in solution without any indication of aggregation or oligomerization (**Supplementary Figure 2**). Furthermore, both DARPin TCEs are highly thermostable, with melting temperatures of 70°C and 60°C for NY_1xCD3 and NY_2xCD3, respectively (**Supplementary Figure 2**). Interestingly, surface plasmon resonance measurements of NY_1xCD3 and NY_2xCD3 binding to HLA-A*0201^+^/NY-ESO1_157-165_(9V) revealed different binding kinetics despite comparable K_D_ values in the single digit nM affinity range (**Supplementary Figure 2**).

The potency and specificity of the DARPin TCE candidates were further assessed in cytotoxic assays using CD8^+^ T cells as effector cells against several different target cells. Similar to the T cell activation screening, both NY_1xCD3 and NY_2xCD3 provoked specific lysis of NY-ESO1_157-165_(9V)-pulsed T2 cells (**Supplementary Figures 3A and 4A**). Additionally, the co-incubation of CD8^+^ T cells with NY-ESO1_157-165_(9C)-positive tumor target cell lines in the presence of either NY_1xCD3 or NY_2xCD3 resulted in efficient and specific killing of antigen-positive, but not antigen-negative tumor cell lines (**Supplementary Figures 3B and 4B**). Finally, both NY_1xCD3 and NY_2xCD3 promoted the efficient killing of NY-ESO1-transduced MCF-7 cells by CD8^+^ T cells (**Supplementary Figures 3C and 4C**).

### High specificity of DARPin TCEs to NY-ESO1_157-165_ with no cross-reactivity to other HLA-A*0201-restricted epitopes

X-scanning mutagenesis, where each peptide residue was mutated to every possible amino acid, indicated, indicated that NY_1xCD3 and NY_2xCD3 interacted with multiple NY-ESO1_157-165_(9V) residues and that few mutations were tolerated at peptide positions p4-p8 and p2-p6 for NY_1xCD3 and NY_2xCD3, respectively (**Figures 1G and 3**). The mutation of peptide residues p4M and p5W, which are both recognized by the TCR 1G4 in ternary complexes,^23^ nearly always abolished recognition by the two DARPin TCEs (**Figure 3**). Moreover, the recognition of HLA-A*0201/NY-ESO1_157-165_ by NY_1xCD3 is mainly governed by the central and C-terminal residues of the peptide (p4-p9), whereas NY_2xCD3 recognition was driven by the central and N-terminal regions (p1-p7). This aligns well with the previous alanine scanning results (**Figure 2C**). Importantly, the HLA-A*0201/NY-ESO1_157-165_-specific DARPins bind across the entire length of the epitope with most substitutions significantly reducing their T-cell activation capacity. As a result, DARPin engagement of even highly similar peptides would most probably be limited or not tolerated.

**Figure 3.**
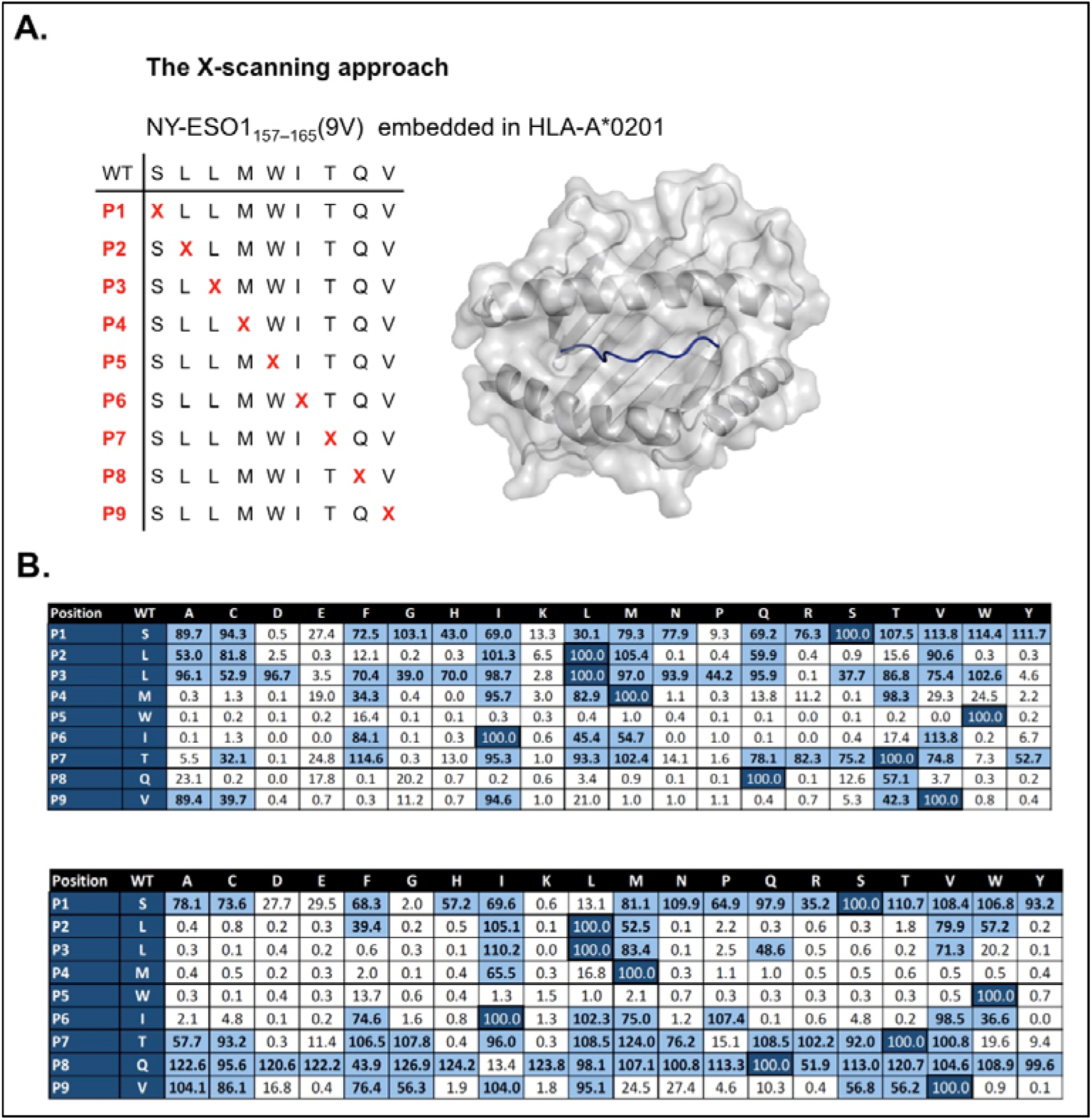
X-scanning mutagenesis demonstrates the high specificity of the DARPin TCEs to HLA-A*0201/NY-ESO1_157-165_ **A.** An X-scanning mutagenesis analysis in which each peptide residue was sequentially replaced by every possible amino acid variant was performed to assess the specificity of DARPins NY_1 and NY_3 to HLA-A*0201/NY-ESO1_157-165_. The peptide NY-ESO1_157-165_ is displayed as a blue ribbon within the peptide binding groove of HLA-A*0201 which surface is colored in transparent grey. **B.** T2 cells were pulsed with each peptide, and thereafter incubated with effector CD8^+^ T-cells and each DARPin TCE at EC_90_ concentrations. The percentage of IFNγ positive CD8^+^ T-cells is presented. Values of independent experiments were averaged and normalized to 100% for the index residue at each position (indicated in dark blue). Values above 30%, indicating significant T-cell activation, are marked in bold and colored light blue. Two independent replicate assays were performed for each DARPin TCE.

To limit the risk of potential cross-reactivity, the X-scanning mutagenesis binding profiles were used to identify, *in silico*, a restricted set of potentially recognized peptide sequences (**Supplementary Table 2**). All identified epitopes were tested for their capacity to mediate CD8^+^ T cell activation against peptide-pulsed T2 cells, in the presence of either NY_1xCD3 or NY_2xCD3. None of the listed epitopes provoked any cross-reactivity with NY_1xCD3, and only two cross-reactive epitopes were identified for NY_2xCD3. Specifically, SLLMWLTPL provoked significant CD8^+^ T cell recognition in the presence of NY_2xCD3, whereas the second epitope TLLIWLFEV induced only weak cross-reactivity (**Supplementary Table 2**).

### Engineered DARPin TCEs with improved efficiency and maintained specificity

We further engineered NY_1xCD3 and NY_2xCD3 to increase their potency without compromising their specificity (**Figure 1H**). First, linkers of different lengths (**Supplementary Table 3**) were inserted between the HLA-A*0201/NY-ESO1_157**-**165_-and the CD3ε-specific DARPin domains in both DARPin TCEs (**Figures 1H and 4A**). These new DARPin TCE constructs were evaluated for their capacity to provoke CD8^+^ T cell activation against either the HLA-A*0201^+^/NY-ESO1_157-165_(9C)-positive cell line IM9 or the HLA-A*0201^+^/NY-ESO1_157-165_(9C)-negative cell line MCF-7 (**Figures 1I and 4A**). Interestingly, DARPin TCE constructs with the shortest, six-amino-acid linkers mediated the most potent CD8^+^ T cell response (**Figures 4A, 4B, and Supplementary Table 3**). Subsequently, we affinity matured CD3ε-specific DARPins and tested the effects on TCE activity of CD3ε-binder variants with affinity values ranging from 6 to 35 nM (**Figure 4C, Supplementary Tables 1 and 3**). Combining the highest affinity CD3ε-specific DARPins paired with the shortest linker mediated the most potent CD8^+^ T cell activation and T cell-mediated cytotoxicity without compromising specificity (**Figures 4C, 4D, and Supplementary Table 3**).

**Figure 4.**
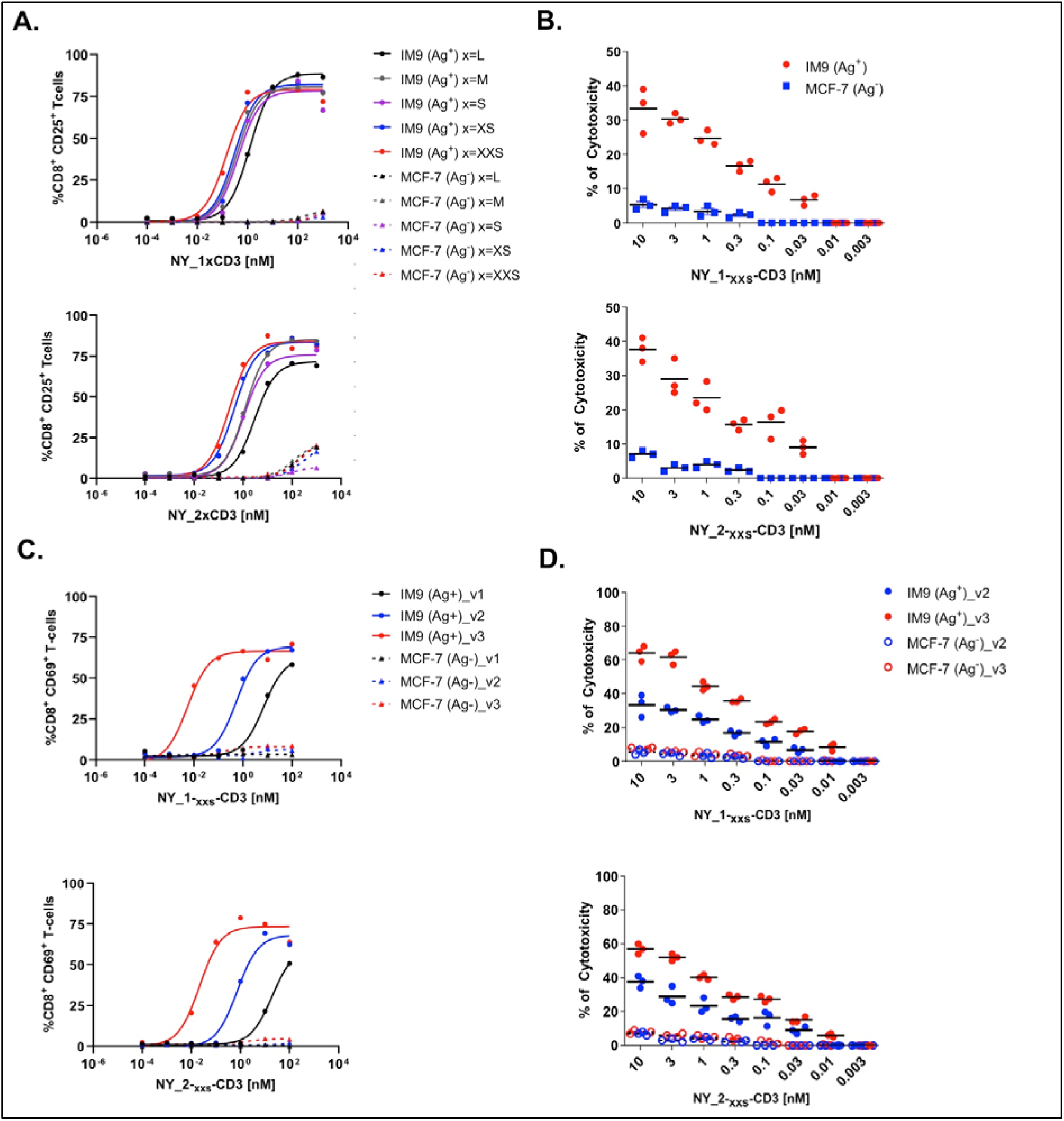
Engineering of linker length and CD3**ε** affinity in DARPin TCEs increases potency without affecting specificity HLA-A*0201^+^/NY-ESO1^+^ (Ag^+^) or HLA-A*0201^+^/NY-ESO1^-^ (Ag^-^) tumor cell lines (IM9 and MCF-7 respectively) were incubated with PBMCs for 48 h in the presence or absence of (**A, B**) NY_1xCD3 or NY_2xCD3 with different linkers (L to XXS), or (**C, D**) sequence optimized versions (v) of NY_1xCD3 or NY_2xCD3 (v1 to v3). The effects on potency of each introduced modification were assessed by measuring the activation markers CD25 (**A**) and CD69 (**C**) on CD8^+^ T cells. The HLA-A*0201^+^/NY-ESO1^+^ or HLA-A*0201^+^/NY-ESO1^-^ tumor cell lines IM9 and MCF-7, respectively, were incubated with effector CD8^+^ T cells in the presence or absence of (**B**) NY_1xCD3 or NY_2xCD3 with the optimal linker length sequence (XXS) or (**D**) the optimized versions (v) of NY_1xCD3 or NY_2xCD3 (v2 and v3). The effects of these modifications were assessed by measuring the percentage of specific lysis obtained on different tumor cell lines using classical chromium release assays. Results are representative of at least two independent experiments.

Finally, we evaluated the most potent lead TCE candidate NY_1xCD3_v3 with a larger panel of tumor cell lines expressing physiological levels of HLA-A*0201^+^/NY-ESO1_157-165_(9C) (**Figures 1I and 5)**. The addition of NY_1xCD3_v3 led to dose-dependent increases of the activation markers CD25 (**Figures 5A and 5B**) and CD69 (**Figure 5C**) on CD8^+^ T cells in the presence of all HLA-A*0201/NY-ESO1_157-165_(9C)-positive tumor target cells but not with antigen-negative cells. Cytotoxic assays performed using the same ensemble of cell lines and CD8^+^ T cells demonstrated that NY_1xCD3_v3 TCE mediated highly efficient and HLA-A*0201/NY-ESO1_157-165_(9C)-specific lysis (**Figures 5D-5F**). In contrast, no significant target cell lysis was observed with NY-ESO1_157-165_(9C)-negative tumor cell lines, even at high DARPin concentrations (**Figures 5D-5F**).

**Figure 5.**
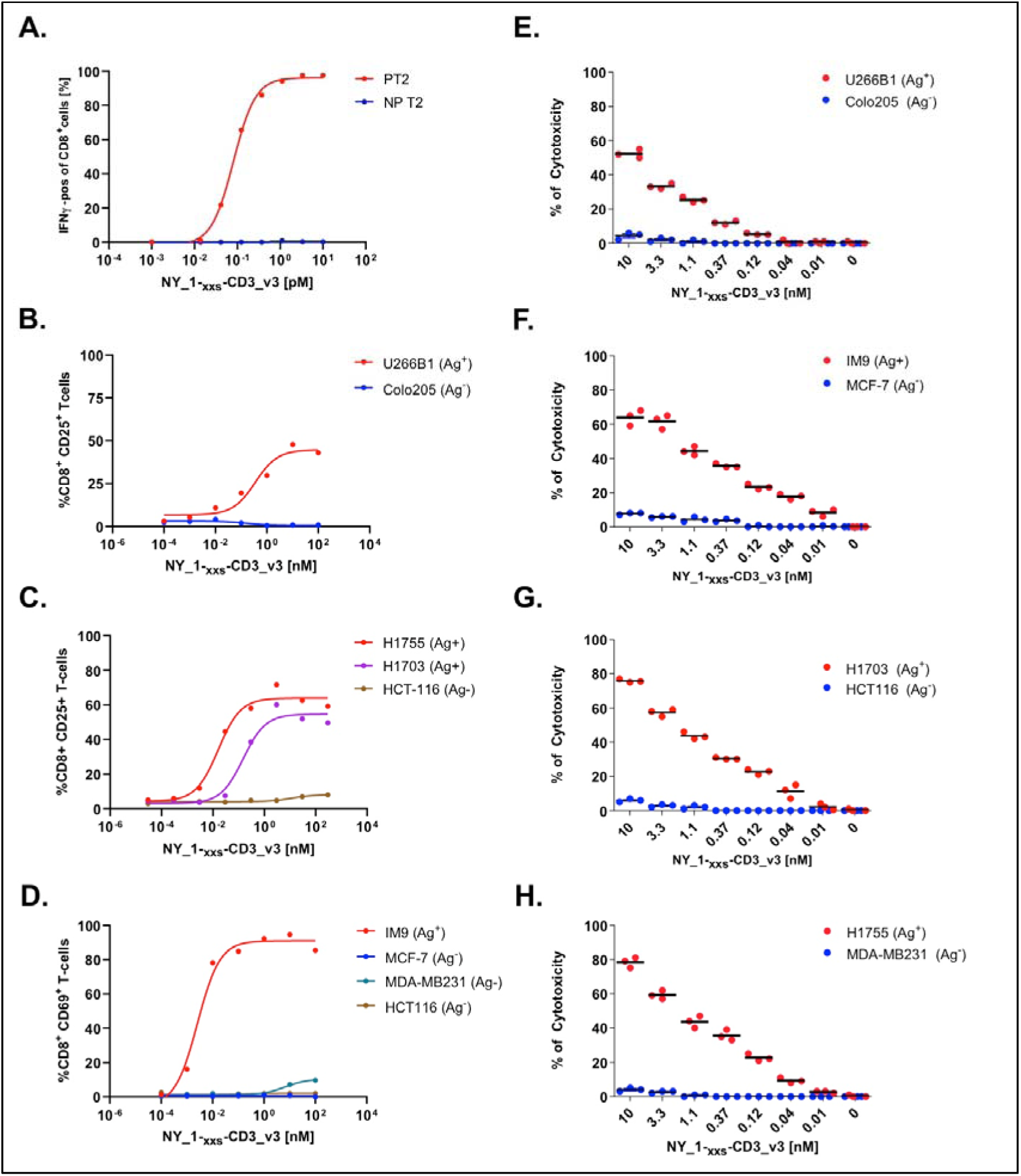
The engineered NY_1xCD3_v3 TCE provokes potent and specific T cell-mediated killing of HLA-A*0201^+^/NY-ESO1_157-165_(9C)^+^ target cells **A.** T2 cells pulsed with 1μM NY-ESO1_157–165_(9V) (PT2) or unpulsed T2 cells (NP T2) were incubated with effector endogenous CD8^+^ T cells in the presence or absence of NY_1xCD3_v3. Intracellular IFNγ levels measured in CD8^+^ T cells are presented. **B**-**H.** The HLA-A*0201^+^/NY-ESO1^+^ (Ag^+^) and HLA-A*0201^+^/NY-ESO1^-^ (Ag^-^) tumor cell lines U266-B1, IM9, NCI-H1755, NCI-H1703 and Colo-205, MCF-7, MDA-MB231 and HCT-116, respectively were incubated with PBMCs (**B**-**D**) or effector CD8^+^ T-cells (**E**-**H**) in the presence or absence of NY_1xCD3_v3. The levels of the activation markers CD25 and CD69 observed in CD8^+^ T-cells are presented (**B**-**D**). The percentage of specific lysis obtained for the different tumor cell lines using chromium release assays are presented in panels **E**-**H**. All results are representative of at least two independent experiments.

### DARPin NY_1 fits snugly to the entire length of the HLA/peptide binding cleft

To assess the molecular bases underlying the affinity and specific binding of NY_1 to HLA-A*0201/NY-ESO1_157-165_(9V), we determined the three-dimensional structure of this ternary complex using single particle cryo-electron microscopy at around 3 Å resolution (**Figures 1J and 6, Supplementary Figure 5, Supplementary Table 4**). Any of the five presented post-processed maps (**Supplementary Figures 6-8**) allowed for unambiguous positioning of the crystal structures of NY_1 (**Supplementary Table 5**) and HLA-A*0201/NY-ESO1_157-165_(9V) ^23^ (**Figures 6A and 6B**). Model-based density-modification ^45^ improved map interpretability notably, yielding a more contiguous density in the DARPin region (**Figure 6D, Supplementary Figures 6-8**). While the DARPin/HLA-peptide interface is well defined in the map, the HLA-A*0201 α3-domain and the solvent-exposed residues of the NY_1 helices are less resolved (**Figure 6A and Supplementary Figures 6-8**). The structures of NY_1 alone and in complex with HLA-A*0201/NY-ESO1_157-165_(9V) were highly similar, with an overall root mean square deviation of 0.4 Å. NY_1 binds on top of HLA-A*0201 with its concave and elongated interaction surface positioned rigidly along the entire length of the peptide-binding cleft. Each constituent helix in the cap elements and internal repeats (IR) is oriented perpendicular to the HLA α1 and α2 helices (**Figures 6A, 6B and 7A**). The N- and- C-Cap elements are positioned above the N- and C-termini of the peptide, respectively, while the IRs of NY_1 engage the central peptide region from residue p4M to p8Q (**Figures 6A and 6D**). The map quality of the IRs is substantially better compared to the Cap elements (**Supplementary Figures 7 and 8**). Remarkably, the distance separating the two broken helices that flank the peptide in HLA-A*0201 appears tailored to closely accommodate the entire width of NY_1 (**Figure 6A**). NY_1 interacts with residues on both heavy chain α-helices and the NY-ESO1_157-165_(9V) peptide (**Figure 6C**). The centrally positioned peptide residues p4M and p5W are surrounded by the hydrophobic NY_1 IR1-residues I46, V48, L53, F56, and the aliphatic part of N-Cap residue R23 (**Figures 6C and 7B**). Additional hydrogen bonds are formed between NY_1 residues IR2-K89 and IR3-Q122 and peptide residues p6I and p8Q, respectively (**Figure 6C**).

**Figure 6.**
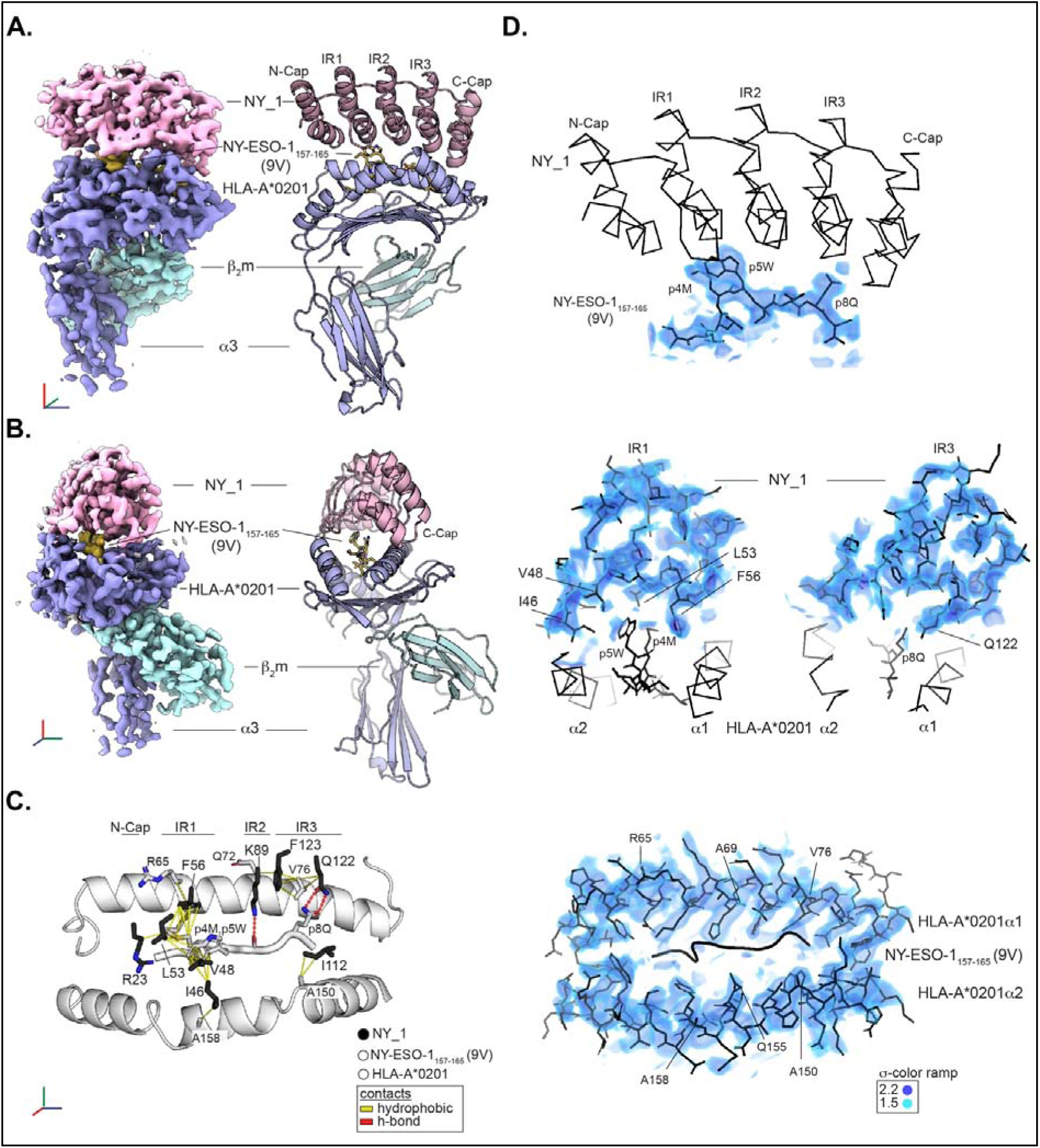
Cryo-EM structure of the NY_1/HLA-A*0201/NY-ESO1_157-165_(9V) complex **A.** Final map and model of the DARPin NY_1 (in light pink) in complex with HLA-A*0201/NY-ESO1_157-165_(9V). Map and model segments of the HLA heavy chain, the β_2_m subunit and the peptide are in light-blue, cyan and yellow, respectively. The Cap and internal repeat (IR) elements of the Darpin NY_1 are indicated. A coordinate frame with blue, green and red axes, indicates the orientation of the complex. **B.** The model and map are represented as in panel A, but shown in a different orientation. **C.** A limited amount of specific close contacts are formed between HLA-A*0201/NY-ESO1_157-165_(9V) and NY_1. The peptide binding cleft of HLA-A*0201 and the presented NY-ESO1_157-165_(9V) epitope are displayed in white. Residues that form interactions with the DARPin NY_1 are shown as sticks. DARPin residues that contact the HLA/peptide complex are shown as black sticks. DARPin residue R23 belongs to the N-Cap element; residues I46, V48, L53 and F56 belong to IR1; K89 belongs to IR2; I112, Q122 and F123 belong to IR3. Hydrophobic contacts and hydrogen bonds are highlighted as yellow and red lines, respectively. **D.** The final map is shown for selected parts of the model overlaid as black lines. The map is presented for the peptide (top), internal repeat elements IR1 and IR3 (middle) as well as MHC helices α1 and α2 (bottom). Visual guides without map are presented as follows: NY_1 as ribbon (top), MHC helices α1 and α2 as well as peptide as ribbon and stick, respectively (middle), and the peptide as cartoon (bottom). Orientations match those of panels A-C. The density is visualized as volume on depicted color ramp. Additional maps, fourier shell correlation curves and map-model cross-correlation are presented in **Supplementary Figures 5-8**.

**Figure 7.**
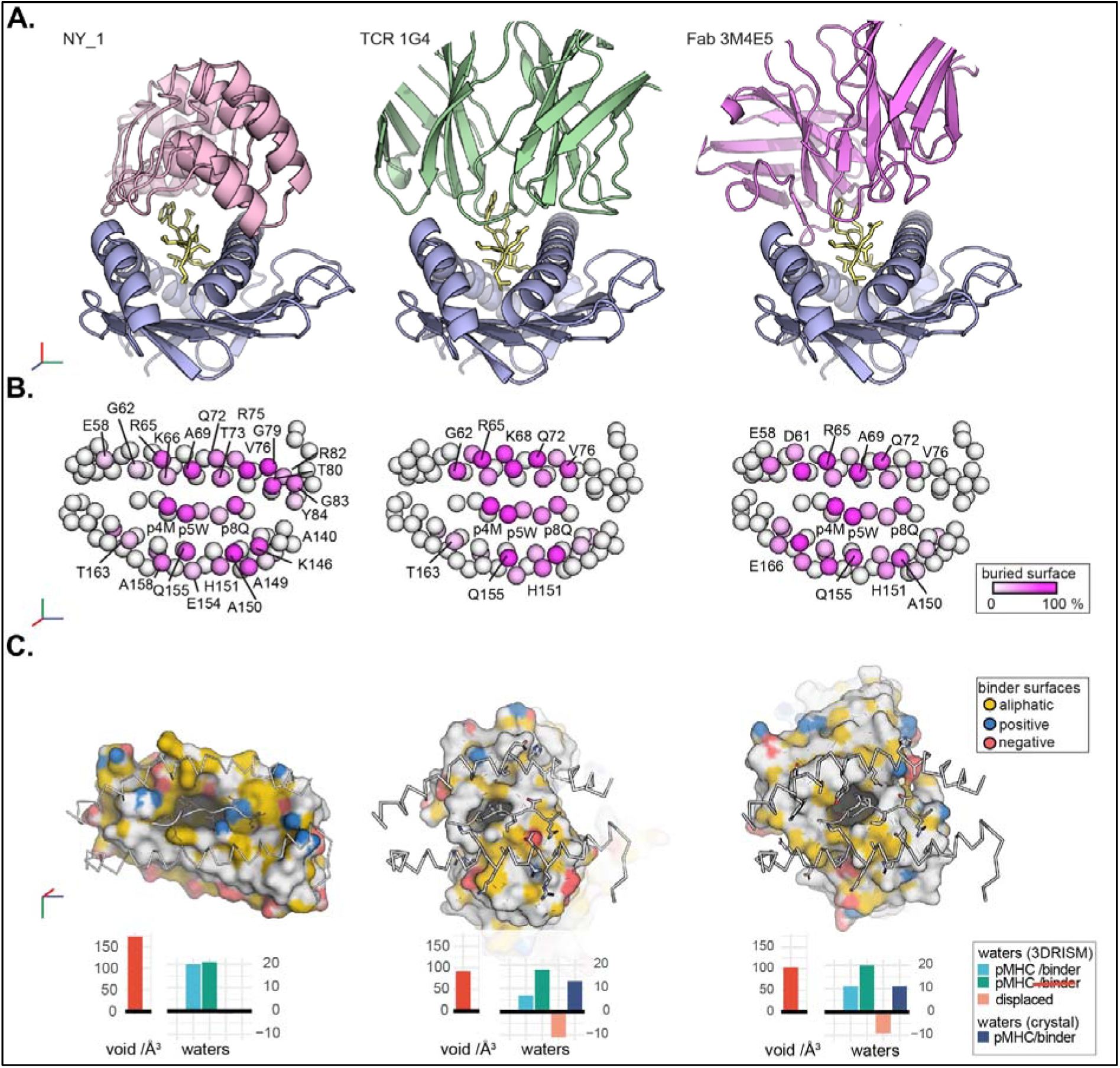
NY_1 fits snugly to the entire length of the peptide binding cleft of HLA-A*0201/NY-ESO1_157-165_(9V) The structure of NY_1/HLA-A*0201/NY-ESO1_157-165_(9V) solved within this study was compared to the previously determined crystal structures of HLA-A*0201/NY-ESO1_157-165_(9V) in complex with the TCR 1G4 or with the Fab fragment 3M4E5. **A.** DARPin NY_1 binds snugly to the entire length of the peptide binding cleft, forming a stiff bridge between the two helices. In contrast, both the TCR 1G4 and the Fab fragment 3M4E5 interact with HLA-A*0201/NY-ESO1_157-165_(9V) using flexible CDR loops that contact the central part of the NY-ESO1_157-165_(9V) epitope and specific HLA-A*0201 residues on both helices. The HLA-A*0201 heavy chain and the peptide are in blue and yellow, respectively. The DARPin, TCR and Fab fragment are in light pink, green and violet, respectively. **B.** The C_α_ atoms of HLA-A*0201/NY-ESO1_157-165_(9V) residues are represented as spheres colored according to buried surface areas upon binding. The white-to-pink color scale is based on the buried surface areas. **C.** Surface views visualize the defined hydrophobic pockets formed by the TCR 1G4 and the Fab 3M4E5 around the key peptide residues p4M and p5W. The top view is rotated by 180° and the atoms of all binders are visualized in yellow, red and blue for hydrocarbon groups without polar substitutions, oxygens of negatively charged and nitrogens of positively charged residues, respectively. The two times larger void present in the NY_1/HLA-A*0201/NY-ESO1_157-165_(9V) interface is predicted to be occupied by a larger number of water molecules (see also **Supplementary Figures 9 and 10**).

We then compared NY_1/HLA-A*0201/NY-ESO1_157-165_(9V) with the ternary complexes formed between the TCR 1G4 or the Fab fragment 3M4E5, and the same HLA/peptide target ^23,46^ to assess similarities and differences in the formed interactions (**Figure 7**). It should be noted that, to our knowledge, all previously determined ternary structures of TCRs in complex with HLA-A*0201/NY-ESO1_157-165_, except for NYE_S3, displayed similar binding interfaces with main contact hot spots focused on peptide residues p4M and p5W.^23,46–49^ The TCR NYE_S3 instead binds an altered conformation of NY-ESO1_157-165_(9V).^47^ A comparative distance-ramp visualization revealed that the complementarity-determining region (CDR) loops of both 1G4 and 3M4E5 form tight interfaces with close binder-to-target distances with the surface of HLA-A*0201/NY-ESO1_157-165_(9V) (**Supplementary Figure 9A**). Despite its distinct scaffold, NY_1 sets a molecular footprint on the HLA-A*0201/NY-ESO1_157-165_(9V) surface that is similar in shape and position to 1G4 bound to the same target, including contacts with the previously defined TCR-restriction triad formed by MHC residues R65, A69 and Q155 ^50,51^ (**Figures 7A and 7B, Supplementary Figure 9**). Strikingly, contacts between the DARPin backbone and HLA-A*0201/NY-ESO1_157-165_(9V) are almost absent, whereas the backbone atoms of both TCRs and Fab fragments contribute to 30-40% of the total binding interface (**Supplementary Figure 9A**). Upon binding of HLA-A*0201/NY-ESO1_157-165_(9V), NY_1 buries an interface area of 1170 Å^2^, which is 10% less than the ∼1300 Å^2^ area buried by 1G4 or 3M4E5 (**Supplementary Figure 9**). All three binders contact both the peptide and the HLA-A*0201 α1 and α2 helices, albeit to differing extents (**Figure 7B and Supplementary Figure 9**). Both NY_1 and 3M4E5 possess larger binding footprints compared to the central diagonal binding mode of 1G4, which form extended interfaces along the α1 and α2 helices (**Figure 7B and Supplementary Figure 9**).

Furthermore, an almost two-fold larger structural void is formed between the DARPin NY_1 and HLA-A*0201/NY-ESO1_157-165_ compared to similar voids found in ternary complexes formed with the TCR 1G4 or the TCR-like antibody 3M4E5. The ∼3 Å resolution structure of the NY_1/HLA-A*0201/NY-ESO1_157-165_ complex did not lead to the detection of any water molecules (**Figure 7C**). However, a solvation prediction algorithm indicated a high probability of the interface between HLA-A*0201/NY-ESO1_157-165_ and NY_1 being filled with more water molecules than the tighter interfaces formed with 1G4 and 3M4E5.

NY_1/HLA-A*0201/NY-ESO1_157-165_(9V) forms a relatively less tight binding interface and creates a solvent-accessible void volume of 175 Å^3^ around the entire length of the peptide (**Figure 7C and Supplementary Figure 10**). In contrast, both 1G4 and 3M4E5 only create distinct hydrophobic pockets around peptide residues p4M and p5W, thus restricting the void volumes to only 100 Å^3^. A solvation prediction algorithm ^52,53^ suggested the larger void formed in the NY_1/HLA-A*0201/NY-ESO1_157-165_(9V) interface to be filled with more water molecules compared to the relatively smaller voids in the interfaces formed with 1G4 and 3M4E5, thus creating a water cushion between NY_1 and HLA-A*0201/NY-ESO1_157-165_(9V) (**Figure 7C, Supplementary Figures 10 and 11**). Coordinated water molecules buried within interfaces play an important role in the fine-tuning of specificity and avidity of protein/ligand interactions,^54–56^ including TCR recognition of HLA/peptide complexes.^57–59^ Furthermore, coordinated water cushions can reduce the entropic costs of ligand binding and significantly increase the affinity between a protein and its ligand.^60^

## Discussion

Developing biological tools, such as soluble TCRs, TCR-like antibodies, and cellular therapeutics, that target low abundance tumor-associated HLA/peptide complexes can be challenging. Common issues include the relative low affinity of these agents, potential cross-reactivity to similar or very different epitopes, as well as the complexity in manufacture and clinical administration. DARPin-based therapeutics offer the potential to overcome these limitations through their excellent biophysical properties and advantageous structural characteristics.^36–38^ When initiating this study, we speculated that the compact and rigid binding domains of DARPins could be particularly well-suited for the size and flat shape of the target surface formed by an HLA/peptide complex. While the flexible CDRs of TCRs or antibodies can allow for cross-reactivity, and thus non-specific binding, the rigid scaffold and binding surface of DARPins typically mediates highly specific interactions with targeted proteins. The present study demonstrates that the developed DARPin molecule NY_1 binds along the entire length of HLA-A*0201/NY-ESO1_157-165_, spanning the distance between the two broken α-helices that flank the peptide binding cleft. Although NY_1 interacts with structural features also bound by TCRs and TCR-like antibodies, the structure of the DARPin molecule allows further contacts with residues at both termini of the HLA-A*0201 peptide binding cleft. Importantly, this enlarged footprint may restrict the capacity of the designed DARPin molecules to cross-react with other HLA/peptide complexes, as demonstrated by the X-scanning mutagenesis assays. Based on these results, NY_1 may display increased specificity to its HLA/peptide target compared to the TCR 1G4 and Fab 3M4E5, due to the combined effects of the absence of mobile and ligand-adaptable CDR loops, a few but specific sidechain-mediated contacts, and a larger water cushion at the interface.

It is well established that multiple mechanisms drive the immunoregulatory function of cancer cells.^61^ Various strategies adopted by cancer cells during tumor progression promote immune escape, including the induction of checkpoint inhibitors or the downregulation of the MHC class I and class II antigen processing pathways. Tumor cells further make use of a large number of immune cells, including regulatory T cells, neutrophils, and macrophages, to help induce an immunosuppressive tumor microenvironment. These tumoral properties, combined with the formation of extracellular matrices and angiogenesis, lead to immune escape, tumor progression, and resistance to therapy.^62–64^ Our strategy in the present study was thus to combine DARPin molecules with high affinity and exquisite specificity towards CD3ε and HLA/TAA complexes, creating TCEs that should be able to recruit CD8^+^ T cells activating and redirecting these for efficient elimination of tumor cells. The functional results demonstrate that the created TCEs promote the efficient killing of a large panel of HLA-A*0201/NY-ESO1_157-165_(9C)-positive tumor cells. In contrast, no killing was observed with NY-ESO1_157-165_(9C)-negative tumor cell lines. However, transfection of the HLA-A*0201^+^/NY-ESO1^-^ cell line MCF-7 with a construct encoding for the full-length NY-ESO1 protein resulted in highly efficient killing by endogenous CD8 T cells in the presence of our TCE constructs. Furthermore, different properties in the linker between the two DARPins and affinity of the CD3ε-specific DARPin altered the efficacy of the TCEs. We hypothesize that a shorter linker between the HLA/peptide- and CD3ε-binding DARPins forces the recruited CD8^+^ T cells within closer proximity of the tumor cells, potentially leading to more efficient synapse formation. Another non-excluding possibility is that the contact half-life between CD8 T cells and tumor target cells is prolonged, allowing extended time for effector cells to kill tumor cells. Finally, the results demonstrate that the higher affinity of the final selection of CD3ε-specific DARPins may elicit significantly stronger synapse formation, leading to higher activation of CD8^+^ T cells. Importantly, this capacity for higher activation did not lead to the unspecific killing of NY-ESO1-negative tumor cells.

A common strategy for developing TCR therapies is to identify HLA/TAA-specific TCRs and thereafter increase their affinity towards the target through directed mutagenesis.^65,66^ Although it has definitively been shown that such therapeutic approaches are often efficacious in both preclinical and clinical assays, the engineered higher TCR affinity often also leads to significantly increased cross-reactivity towards other unwanted targets.^67,68^ Furthermore, a previous study revealed that a chimeric antigen receptor based on the antibody 3M4E5 caused moderate lysis of HLA-A*0201-expressing targets due to enhanced T cell avidity, independent of the presented antigen and despite the high affinity of the Fab fragment 3M4E5 to HLA-A*0201/NY-ESO1_157-165_. It was, therefore, required to to reduce the affinity of the TCR-like antibody through structure-guided mutagenesis to levels equivalent to the conventional TCR to ensure specificity.^69^ In contrast, we demonstrated that high-affinity DARPin TCEs, selected from naïve libraries in four rounds of *in vitro* selection, still maintain very high specificity.

In conclusion, DARPins offer a promising foundation for developing HLA/peptide-specific therapeutics with enhanced potency. Our presented workflow allows for rapid identification and isolation of lead DARPin candidates towards both HLA class I and class II molecules in complex with TAAs, as well as pathogen-derived epitopes. As exemplified by our creation of bispecific DARPin TCEs, tumor or virus-specific DARPins could be easily combined with any of the CD3ε-specific DARPins.

## Material and methods

### Production of biotinylated MHC/peptide complexes for the selection of DARPin candidates

Complexes of HLA-A*0201 with human β_2_-microglobulin (hβ_2_m) and each of the peptides NY-ESO1_157-165_(9V) (SLLMWITQV), NY-ESO1_157-165_(9VAA) (SLL<u>AA</u>ITQV) and EBNA-1 (FMVFLQTHI), were produced as previously described.^70,71^ A codon-optimized HLA-A*0201 (HLA-A*0201avi) construct that comprised a linker (GSGGSGGSAGG) and an avi-biotinylation tag (GLNDIFEAQKIEWHE) was used for protein expression in *E. coli* BL21 (DE3) at 37°C as inclusion bodies (IB). The IB protein contents were thereafter purified and dissolved in 50 mM MES, 5 mM EDTA, 5 mM DTT, 8 M urea pH 6.5. HLA-A*0201avi (or HLA-A*0101avi) and hβ_2_m were refolded with each peptide using final concentrations of 25, 30 and 15 mg, respectively, per 500 mL volume in 50 mM Tris pH 8.3, 230 mM L-arginine, 3 mM EDTA, 255 μM GSSG; 2.5 mM GSH and 250 μM PMSF. For biotinylation, each refolded complex was concentrated to a volume of 7.5 mL, and buffer was exchanged to 100 mM Tris pH 7.5, 150 mM NaCl, 5 mM MgCl2 pH 7.5 using PD10 columns. Avi-tagged heavy chains were biotinylated by adding 5 mM ATP, 400 µM Biotin, 200 µM PMSF and 20 µg BirA enzyme. Each refolded biotinylated complex was isolated using size exclusion chromatography (Superdex 200 HiLoad 16/600, Cytiva) in PBS supplemented with 150 mM NaCl, 1 mM EDTA, 10% glycerol. Samples were concentrated to about 1 mg/mL and flash-frozen as 25 and 50 µl aliquots using liquid nitrogen.

### Ribosome display selection of HLA-A*0201/NY-ESO1_157-165_(9V)-specific DARPin candidates

Four DARPin libraries (N2C and N3C) were used in ribosome display selections ^35,72,73^ against the refolded biotinylated HLA/peptide complexes. Four selection rounds were performed for each pool. To direct DARPin binding towards the peptide and not only the HLA-A heavy chain scaffold, deselection steps were performed using the complexes HLA-A*0201/NY-ESO1_157-165_-9VAA and HLA-A*0201/EBNA-1. Selection stringency was continuously increased to enrich high affinity binders. Nunc MaxiSorp 96-well microplates (Thermo Fisher Scientific, Zurich, Switzerland), coated with 100 µl solution of 66 nM neutravidin in PBS and incubated at 4°C overnight, were used for panning. Microplates were washed three times the following day with 300 µl PBST per well and blocked with 300 µl PBST-BSA for 1 h at 4°C, rotating at 700 rpm, prior to de-selection. After emptying the wells, 100 µl of 50 nM biotinylated HLA/peptide deselection target in PBST-BSA (PBS pH 7.4, 0.05% Tween 20, 0.2% (w/v) BSA) was added to each well, rotating at 700 rpm at 4°C for 1 h. During this incubation step, *in vitro* mRNA translations were performed. Shortly after translations and generation of ternary ribosomal display complex mixes, solutions were discarded and microplate wells were washed three times with 300 µl PBST and finally incubated with a washing buffer with tween (WBT) containing 0.2% (w/v) BSA: 50mM Tris-HOAc (pH 7.5 at 4°C), 150 mM NaCl, 50 mM Mg(OAc)_2_, 0.05% Tween 20. For each de-selection step, WBT-BSA solutions were discarded and aliquots (150 µl for first selection round and 100 µl for rounds 2-4) of the translated ternary complexes were transferred subsequently three times to a prepared Nunc MaxiSorp well containing immobilized deselection HLA/peptide targets, and incubated for 20 min at 4°C. At the end of the deselection process, ternary ribosomal display complexes were used in the selection step on HLA-A*0201/NY-ESO1_157-165_(9V).

### Binding screening of DARPins to HLA-A*0201/NY-ESO1_157-165_(9V) using homogenous time-resolved FRET (HTRF)

DARPin clones selected by ribosome display were cloned into a derivative of the pQE30 (Qiagen) expression vector and transformed into *E. coli* XL1-Blue (Stratagene), plated out on LB agar/ampicillin and incubated overnight at 37°C. Single colonies were picked into individual wells of twelve 96-well plates containing 165 μl growth medium (LB containing 1% glucose and 50 μg/ml ampicillin) and incubated overnight (37°C/800 rpm). 150 μl of fresh LB medium containing 50 μg/ml ampicillin was inoculated with 8.5 μl of the overnight culture for expression induced with isopropyl β-D-1-thiogalactopyranoside (IPTG) (0.5 mM for 6 h). Cells were harvested by centrifugation of 96-deep-well plates before resuspension in 8.5 μl B-PERII (Thermo Scientific) and incubation for 1 h at RT/600 rpm. 160 μl PBS was added and cell debris removed by centrifugation. The extract was diluted 1:200 in PBSTB (PBS, 0.1% Tween 20, 0.2% [w/v] BSA, pH 7.4) with 20 nM biotinylated target, 1:400 anti-6His-D2 HTRF antibody (Cisbio International, Gif-sur-Yvette, France) and 1:400 anti-strep-Tb (Cisbio, France) in a 384-well format (120 min incubation at 4°C). Plates were read with a Tecan M1000 using standard HTRF settings. Extracts of each lysed clones were tested for binding to biotinylated HLA-A*0201/NY-ESO1_157-165_(9V) and to the negative controls HLA-A*0201/NY-ESO1_157-165_(9VAA) and HLA-A*0201/EBNA-1. The specificity of each identified DARPin clone to HLA-A*0201/NY-ESO1_157-165_(9V) was assessed by calculating the ratio of the obtained HTRF signal against the HTRF signal for the two HLA/peptide negative controls. All DARPin binders which generated 25-times higher HTRF signals from their interactions with HLA-A*0201/NY-ESO1_157-165_(9V) *versus* the negative controls were considered as specific preliminary hits and were taken forward for subsequent characterizations.

### Production of DARPin candidates

The 26 best DARPin candidates, identified in the initial screening for HLA-A*0201/NY-ESO1_157-165_(9V), were purified using immobilized metal affinity chromatography (IMAC) in 96-well format and thereafter re-buffered in PBS. A single colony was picked for each DARPin into TB medium (containing 1% glucose and 50 μg/ml ampicillin) and incubated overnight at 37°C for 16 h, shaking at 700 rpm. Fresh TB medium (50 μg/ml ampicillin) was inoculated with the overnight culture at a ratio of 1:10 and incubated at 37°C at 700 rpm. After 2 h, protein expression was induced by the addition of IPTG (0.5mM), and bacterial cultures were kept for 6 h at 37°C at 700 rpm. Cells were harvested by centrifugation (3200 g for 6 min) and thereafter lysed by incubation with B-PER II (Bacterial Protein Extract Reagent, Thermo Fisher), DNAse I (200 U/ml), and lysozyme (0.4mg/ml) for 60min at room temperature and 900 rpm. Cell debris was removed by centrifugation at 3200 g for 60 min at 4°C. The 0.65 µm filtered supernatant was purified by IMAC (HisPur Cobalt Spin Plates, Thermo Fisher) via the N-terminal His-tag, including washing steps and subsequent elution with 150 mM imidazole. Finally, purified DARPin molecules were re-buffered into PBS using Zeba^Tm^ Spin Desalting Plates (Thermo Scientific).

### Surface plasmon resonance assessment of binding of DARPin candidates to HLA/peptide targets

An inverse ProteOn setup was used to preliminarily test the interactions of each identified DARPin candidate to HLA-A*0201/NY-ESO1_157-165_(9V). An RGS-His antibody (Qiagen ID: 34650) was immobilized on a GLC chip, allowing the capture of the MRGS-6His tagged DARPin molecules and their use as ligands. HLA-A*0201/peptide targets were immobilized as analytes on a neutravidin chip at different dilutions in a tris-based WBT buffer or a HEPES buffer.

### Design and optimization of DARPin T-cell engagers using a selection of anti-CD3**ε−**specific DARPins

Three different variants for the anti-CD3ε-specific DARPin domain of the bispecific T-cell engager were investigated. The three variants resulted from affinity maturations (data not shown) and were used to evaluate the best-suited affinity for T-cell engagement. Sequences and point mutations incorporated in the three DARPin variants are described in **Supplementary Table 1**.

### Design of linkers for optimization of the bispecific DARPin T-cell engagers

The importance of various peptide linkers with different lengths and compositions for the highest possible CD8^+^ T cell engagement while maintaining specificity was evaluated following their introduction between HLA-A*0201/NY-ESO1_157-165_(9V)-binding DARPins and a selection of CD3ε-binding DARPins. Five different linker lengths were tested: XXS (GSPTGS), XS (GSPTPTPTTGS), S (GSPTPTPTTPTPTPTTGS), M (GSPTPTPTTPTPTPTTPTPTPTGS), and L (GSPTPTPTTPTPTPTTPTPTPTTPTPTPTTPTPTPTGS).

### Size Exclusion Chromatography (SEC) analyses

DARPin molecules were run on an analytical size exclusion chromatography column (Superdex 200, GE Healthcare) at 2 mg/ml in PBS buffer and evaluated for monodispersity, aggregation, and oligomerization.

### Circular dichroism measurements

Circular dichroism measurements were performed with a Jasco J-815 using a 1 cm pathlength cuvette (Hellma). The molar residue ellipticity (MRE) at 222 nm was measured over a temperature ramp from 20°C to 90°C. Spectra from 190-250 nm were taken before and after the variable temperature measurement at 20°C. Samples were measured at 1μM in PBS.

### Alanine scanning mutagenesis assays

Each NY-ESO1_157-165_(9V) residue in was sequentially replaced by alanine in the indicated position. Anchoring positions p2 and p9 were substituted simultaneously. T2 cells pulsed with 10^-6^ M of each mutated peptide version were incubated with effector CD8^+^ T cells for 4 h in the presence of DARPin candidate concentrations allowing EC_90_ levels for NY-ESO1_157-165_. Intracellular IFNγ on CD8^+^ T cells was detected by FACS using appropriate antibody (BD, 554702).

### **X-** scanning mutagenesis assays

Each peptide position in NY-ESO1_157-165_(9V) was mutated to each of the 19 other amino acids. T2 cells were pulsed with each of the mutated peptides and incubated with effector CD8^+^ T cells for 4 h in the presence of DARPin candidate at a concentration allowing EC90 levels of T cell activation in the presence of the wild-type peptide. Intracellular IFNγ on CD8^+^ T cells (BD, 554702) was detected by FACS. Each of the experiments were performed in two independent replicates. Values were averaged and normalized to 100% for the according wild-type residue in each position. A threshold of 30% T-cell activation was used to identify potential amino acid substitutions in each position of the NY-ESO1 peptide for the two candidates NY-1 and NY-2. With this information, the according peptide motifs [ACFGHILMNQRSTVWY]-[ACILMQV]-[ACDFGHILMNPQSTVW]-[FILMT]-W-[FILMV]-[CFILMQRSTVY]-[QT]-[ACITV] and [ACFHIMNPQRSTVWY]-[FILMVW]-[ILMQV]-[IM]-W-[FILMPVW]-[ACFGILMNQRSTV]-[ACDEFGHKLMNPQRSTVWY]-[ACFGILSTV] for NY-1 and NY-2, respectively, were entered into the ScanProSite to scan for peptides matching this motif in the SwissProt database on the Expasy site (https://prosite.expasy.org/scanprosite/; Option 2 - Submit MOTIFS to scan them against a PROTEIN sequence database; filtered for homo sapiens taxonomy). Identified potentially cross-reactive peptides for NY-1 and NY-2 (**Supplementary Table 2**) were ordered at Genscript (Piscataway, NJ, USA) and used to evaluate the capacity of the DARPin T cell engagers to mediate activation of CD8^+^ T cells in the presence of pulsed T2 cells.

### CD8^+^ T cell preparation

Blood samples from anonymous healthy volunteers were purchased from the Karolinska University Hospital. Peripheral blood leukocytes were isolated by Ficoll density gradient centrifugation. CD8^+^ T cells were thereafter isolated using well-established protocols (Milteny Biotec, Germany). Purity was checked by flow cytometry and cells were thereafter seeded in RPMI 1640 + 10% FBS, and 1 μg/ml of phytohaemagglutinin (PHA; Thermofisher) at 10^6^ cells/ml. Cells, incubated at 37°C for three days, were then collected, washed and 20 ng/ml of human IL-2 (Immunotools) was added.

Cells were kept for 4-6 days at a density of 10^6^ cells/ml in a 6-well plate.

### T cell activation assays using T2 cells

T2 cells were pulsed overnight at 37°C with 10 μM of the NY-ESO1_157-165_(9V) peptide in serum free medium. Unpulsed T2 cells were used as control. Effector CD8^+^ T cells were added at an effector to target ratio 1:5 in the presence of DARPins at the indicated concentrations and incubated for 4-5 hours at 37°C. Intracellular IFNγ (BD, 554702) was detected by FACS.

### T cell activation assays using tumor cell lines

HLA-A*0201^+^/NY-ESO1^+^ and HLA-A*0201^+^/NY-ESO1^-^ tumor cells were incubated with PBMCs at an effector-to-target ratio of 10:1 for 48 h at 37°C. Levels of activation were followed by assessment of the expression levels of the surface markers CD25 and CD69 on CD8^+^ T cells by FACS. Supernatants were collected and IFNγ quantification was performed using the Luminex technology. Antibodies used for FACS analysis are listed in **Supplementary Table 6**.

### Chromium release cytotoxicity assays

The efficiency of DARPins to kill target cells was tested in standard chromium 4 h (^51^Cr, Perkin-Elmer) release assays using triplicate cultures in round bottom 96-well plates. Briefly, HLA-A*0201^+^/NY-ESO1^+^ or HLA-A*0201^+^/NY-ESO1^-^ target cells, or peptide-pulsed versus unpulsed T2 cells were prepared by incubating them with 10 ml of 10 mCi/ml ^51^Cr for 1 h, washed, seeded (1000 cells in 50 μl of complete T cell medium) and thereafter incubated for 4 h with a 10:1 ratio of CD8^+^ T cells in the presence of different concentrations of DARPin molecules. Chromium release was measured in the harvested supernatants after 4 h incubation at 37°C. The percentage of specific lysis was calculated according to the formula: % specific lysis = ((experimental release - spontaneous release)/(maximum release - spontaneous release)) x 100.

### Cell lines

The cell lines IM-9, NCI-H1755, NCI-H1703, and U266 were used as HLA-A*0201^+^/NY-ESO1^+^ cells. Cell lines MDA-MB231, HCT116, Colo-205 and MCF7 were used as HLA-A*0201^+^/NY-ESO1^-^ cells. All cells were purchased from the American Type Culture Collection. MCF-7 transduced with NY-ESO1 were generated in house. HCT116 were cultured in McCOY’s 5a (Gibco), and T2, U266, Colo205 and IM-9 were cultured in RPMI 10 medium (Gibco). The rest of the cells were maintained in DMEM (Lonza) medium. All different medium were supplemented with 10% heat-inactivated FCS (fetal calf serum, Biochrom), 2 mM L-glutamine, 1 mM HEPES and 100 U/ml penicillin/streptomycin. All cell culture flasks and plates were purchased from BD Falcon (Franklin Lakes, NJ). To pulse T2 cells, 10μM of the NY-ESO1_157-165_(9V) peptide was added in RPMI medium (serum free) for 16 h at 37°C. Unpulsed cells were used as control. All cell lines (**Supplementary Table 7**) were tested negative for mycoplasma.

### Crystal structure of NY_1

While screening for crystals of the NY_1/HLA-A*0201/NY-ESO1_157-165_(9V) complex, concentrated to ∼10 mg/mL in 20 mM HEPES, 10% glycerol, 300 mM NaCl pH 7.5, we obtained a dataset of NY_1 alone at a resolution of 1.8 Å. Crystallization was performed within the protein science facility at Karolinska Institutet (https://ki.se/en/mbb/psf-mx). Crystals of NY_1 alone were obtained in condition 1-48 (pH 8.5; 0.12 M alcohol mix; 0.1 M buffer system 3; 37.5% (v/v) MPD_P1K_P3350) of the Morpheus screen (Molecular Dimensions Ltd.). For condition screening, the Mosquito robot was used to setup 1:2, 1:1 and 2:1 drop ratios in 96-well sitting-drop iQ TTP Labtech plates (TTP Labtech). The dataset was collected at a wavelength of 0.96880 Å at the I24 microfocus beamline at the Diamond synchrotron light source (DLS, UK). The structure was solved by molecular replacement, using the previously determined crystal structure of another DARPin molecule (PDB code 2XEH ^74^). The model was re-built and refined iteratively using Coot ^75^ and Phenix ^76^ to final R and R_free_ values of 18.3 and 21.5%, respectively. The structure was deposited under PDB code 9EPA (**Supplementary Table 5**).

### Cryo-electron sample preparation and data acquisition

The Avi-tag of HLA-A*0201 was substituted by Strep-tag-II (STII) applying sequence and ligation independent cloning (SLIC).^77^ SLIC was also applied to introduce a Tobacco Etch Virus protease (TEV) cleavage site following the N-terminal His tag of DARPin NY_1. Refolded HLA-A*0201-STII/NY-ESO1_157-165_(9V) was obtained as described above but purified further through Strep-Tactin Superflow high capacity columns (1 mL, IBA lifescience) run in 20 mM HEPES, 300 mM NaCl pH 7.5. After a column wash, the protein was eluted using the same buffer supplemented with 2.5 mM desthiobiotin. TEV-cleaved DARPin NY_1 was reverse-purified via IMAC and the monomer was isolated from Superdex 200 equilibrated in 20 mM HEPES, 150 mM NaCl pH 7.5. For cross-linking, HLA-A*0201/NY-ESO1_157-165_(9V) was mixed in a 1:2 molar ratio with NY_1, concentrated to an absorbance at 280 nm (Abs280) of 2.3, and incubated for 45 min in 25 mM HEPES, 150 mM NaCl, supplemented with 1.4 mM BS(PEG)5 (BS5) (Thermo Fisher Scientific). The cross-linker was quenched by addition of 25 mM Tris. Samples for grid screening were isolated from Superdex 200 GL 10/300 SEC in 25 mM HEPES, 150 mM NaCl, pH 7.4, and concentrated to Abs280 values of ≥ 2 (**Supplementary Figure 12**).

UltrAuFoil holey grids (R 0.6/1 gold foil on gold 300 mesh; Quantifoil Micro Tools GmbH) were glow-discharged for 60 s at 20 mA using a Glo-Qube (Quorum) instrument. Purified NY_1/HLA-A*0201/NY-ESO1_157-165_(9V) complexes were thawed, centrifuged (14,000 g for 5 min at 4°C) and diluted to ∼0.5 mg/ml with gel filtration buffer. Protein was loaded into the freshly glow-discharged grids and plunge-frozen in LN_2_-cooled liquid ethane using a Vitrobot Mark IV (Thermo Fisher Scientific) with a blot force of 0 for 4 s. Temperature and relative humidity were maintained at 4°C and 100%, respectively. Grids were clipped and loaded into a 300-kV Titan Krios G3i microscope (Thermo Fisher Scientific, EPU 2.8.1 software), equipped with a Gatan BioQuantum energy filter and a K3 Summit direct electron detector (AMETEK). Grids were screened for quality control based on particle distribution and density, and images from the best grid were recorded. Micrographs were recorded at a nominal magnification of ×130,000, corresponding to a calibrated pixel size of 0.648Å. The dose rate was 10.4 electron physical pixels per second, and images were recorded for 2.3 s divided into 45 frames, corresponding to a total dose of 57.5 electrons per Å^2^. Defocus range was set between -0.6 μm and -2.5 μm. Gain-corrected image data were acquired.

### Cryo-EM data processing

The data processing workflow for the NY_1/HLA-A*0201/NY-ESO1_157-165_(9V) dataset is illustrated in **Supplementary Figure 5**. Image processing steps were performed with cryoSPARC v4.0.3.^78^ Movie stacks were motion-corrected and contraction transfer function (CTF) estimated within the CryoSPARC Live Session, using Patch Motion Correction and Patch CTF estimation. Poor-quality micrographs were discarded based on an upper CTF fit threshold of 10Å, resulting in 7,781 micrographs used for further processing. Picking of particles was initiated during CryoSPARC Live session with blob-picker, and live 2D classification of approximatively 100,000 particles yielded five ‘protein-like’ 2D classes of 12,000 particles that were used as templates for further live template picking, yielding approximatively 7.7 million particles. A selection of 2D classes comprising 100,000 particles were used for *ab initio* reconstruction without imposing symmetry (C1) to generate two initial models from which the most protein-like model was chosen. Repeated 2D classification further cleaned the particles down to a selection of 205,000, which were classified into three classes with heterogenous 3D refinement resulting in three seemingly identical classes. All 205,000 particles were used for homogenous 3D refinement, local CTF refinement followed by non-homogenous refinement ^79^ (**Supplementary Figure 5**). Local resolution was estimated showing little deviation in resolution from the overall resolution of 3.0 Å. 3DFSC analysis revealed anisotropy along the Y-direction but confirmed the resolution with an anisotropy-corrected global estimate of 3.1 Å. Auto-sharpening and density-modification in Phenix improved the map significantly (**Supplementary Figures 6-8**). Post-processing using DeepEMhancer ^80^ did not improve the map in the relevant region of the DARPin/HLA-peptide interface.

### Cryo-EM model building and refinement

The crystal structures of NY_1 and HLA-A*0201/NY-ESO1 (PDB 1S9W ^81^) were placed into the initial map in ChimeraX ^82^. The model was iteratively refined by model building in Coot and ChimeraX-Isolde, as well as Phenix real space refinement with integrated amber force field with Ramachandran, secondary structure and reference structure restraints.^75,83–85^ Post-processed maps were obtained in cryoSPARC and Phenix, respectively.^45,80,86^ Progress of structure refinement and map quality improvements were validated by Fourier-Shell correlation plots, map-model CC, EM-ringer score and model geometry statistics as available in Phenix ^83^ and Molprobity ^87^. The density modification algorithms had the largest impact on the DARPin map, while the quality of the auto-sharpened HLA/peptide map was almost identical to its density-modified version.

### Structural analysis and visualization

Structures were analyzed and visualized using ChimeraX ^82^ and PyMOL (Schrödinger). Molecular interactions and buried surface areas were analyzed applying Arpeggio ^88^ and PISA ^89^ webservers. Void volumes and the predicted numbers of water molecules were obtained from CastP ^90^ and Amber 3DRISM analyses.^52,53^ Analyses were tidied and visualized using custom-made scripts applying mainly the R tidyverse and ggplot2 packages.

## Data availability statement

The data supporting the findings of this study are available from the corresponding author upon reasonable request.

## Acknowledgements

Parts of the computations were performed at the NSC Tetralith provided by the National Academic Infrastructure for Supercomputing in Sweden (NAISS) and PReSTO funded by the Swedish Research Council through grant agreement no. 2022-06725 (NAISS) and no. 2018-06479 (PReSTO). Parts of this work were facilitated by the Protein Science Facility at Karolinska Institutet (http://ki.se/psf)??? where we would like to thank M. Moche for assistance. We would like to thank Diamond Light Source for beamtime and the staff of beamline I24 for assistance with crystal testing and data collection. Cryo-EM data was collected at the Cryo-EM Swedish National Facility funded by the Knut and Alice Wallenberg, Family Erling Persson and Kempe Foundations, SciLifeLab, Stockholm University and Umeå University.

## Author contributions

AA, VL and TS conceptualized the initial theoretical approach. AA, VL, TS, NV, MW, TSch, KW, SM and MC designed research and analyzed data. NV, TSch, SM, KW, SF, NK, TR, MP, NP, FR, DV, SB, AC, TH, XH, RS, EA, HGL, BJC, MC, TS and MW generated and/or interpreted research results. AA wrote the first draft of the manuscript, and NV, TS, TR, SM, KW, MC, TSch, BJC, EA and HGL contributed to the finalization of the manuscript.

