## Supplementary Information for "Development of DARPin T cell engagers for specific targeting of tumor-associated HLA/peptide complexes"

**Supplementary Tables and Figures**

**Supplementary Table 1. Amino acid sequences of the three CD3ε-specific DARPin variants used within this study**

| **CD3ε-specific DARPins** | **AA sequence** (N-cap/ IR1/ IR2/ C-Cap) |
| --- | --- |
| version 1  K_D_: 35 ± 21 nM | DLGQKLLEAAWAGQDDEVRELLKAGADVNA  KNSRGWTPLHTAAQTGHLEIFEVLLKAGADVNA  KDDKGVTPLHLAAALGHLEIVEVLLKAGADVNA  QDSWGTTPADLAAKYGHEDIAEVLQKAA |
| version 2  K_D_: 14 ± 3 nM | DLGQKLLEAAWAGQDDEVRELLKAGADVNA  KNSRGWTPLHTAAQTGHLEIFEVLLKAGADVNA  K**N**DK**R**VTPLHLAAALGHLEIVEVLLKAGADVNA  **R**DSWGTTPADLAAKYGH**Q**DIAEVLQKAA |
| version 3  K_D_: 6 ± 0 nM | DLGQKLLEAAWAGQ**L**DEVR**I**LLKAGADVNA  KNSRGWTPLHTAAQTGHLEIFEVLLKAGADVNA  K**TN**K**R**VTPLHLAAALGHLEIVEVLLKAGADVNA  **R**DTWGTTPADLAAKYGH**R**DIAEVLQKAA |

DARPin amino acid sequences for the four repeats of CD3ε-specific DARPin versions 1-3.

Listed K_D_-values were obtained by surface plasmon resonance.

Mutations identified during affinity maturation rounds are in red.

**Supplementary Table 2. Cross reactive peptides identified through X-scanning analyses of NY_1xCD3 and NY_2xCD3**

| **NY_1xCD3** | | | | **NY_2xCD3** | | | |
| --- | --- | --- | --- | --- | --- | --- | --- |
| **#** | **Sequence /%TCA** | **#** |  | **#** | **Sequence /%TCA)** | **#** | **Sequence /%TCA** |
| 01  02  03  04  05  06  07  08  09  10  11  12  13  14  15  16  17  18  19  20  21  22  23  24  25  26  27  28  29  30  31  32  33  34  35  36  37  38  39  40 | **SLLMWITQC /44%**  AAGTWVLQA  ACGIWMITV  AMLLWVQQA  AQLLWFLQT  AVFLWLVTI  AVLTWLSQT  CVATWVFTA  FIDMWFSTV  FLTLWLTQV  GVTTWIQTV  HCLFWLLQV  HIGTWFTTT  HISTWLYQA  ICLIWLLTV  ICVIWLYTA  LIQFWMSTV  LLGTWVFQV  LLPLWLSTT  LVITWIMTV  NLFLWLSTV  NLLIWVTTI  QISLWIITA  QLSMWIRTC  QMPLWVRQI  QQAMWMMTA  RLCIWLLQT  RVMLWVTTA  SLFLWIRTA  SLSTWIVTV  SQCMWLMQA  TIHTWIRQC  TLGLWLTTA  TLGLWMVTA  TLLLWLCQA  TLPIWMMQT  TLQIWLRQA  TLQTWLVQA  VADTWVLTA  VAPLWMRQI | 41  42  43  44  45  46  47  48  49  50  51  52  53  54  55  56  57  58  59  60  61  62  63  64  65  66  67  68  69  70  71  72  73  74  75  76  77  78  79  80 | VLLLWLLQV  WVSFWISQA  YLNMWITTC | 01  02  03  04  05  06  07  08  09  10  11  12  13  14  15  16  17  18  19  20  21  22  23  24  25  26  27  28  29  30  31  32  33  34  35  36  37  38  39  40 | **SLLMWITQC /63%**  **SLLMWLTPL /63%**  **TLLIWLFEV /7%**  AIVIWFTGF  AIVIWIILA  ALVIWWQRV  AMVIWINEI  CFIMWVLFI  CIIIWLLAG  CIVMWLAGG  CLLIWLLDA  CLMIWLIFS  CMVIWVLAF  CVLIWVVGG  FLIIWLITG  FLVIWILFS  FLVIWLVGF  HLVIWLLLV  IFLIWLLDF  ILMIWLMAT  IMIIWVLAI  IVIIWIVSC  IVLIWVIAC  IVLIWVIAC  IVLIWVVSV  IVVIWVIVS  MFIMWFSGL  MLVIWILTL  NIQIWLANG  NIVIWVSGS  NLLIWVTTI  NLVIWPSVA  NVLIWPMEG  NVLIWPTDG  PLLMWLLKS  PLMIWVTDT  PLVMWLQGG  PVLMWVQAL  RLIIWILYL  RLQIWPGYA | 41  42  43  44  45  46  47  48  49  50  51  52  53  54  55  56  57  58  59  60  61  62  63  64  65  66  67  68  69  70  71  72  73  74  75  76  77  78  79  80 | RLVIWPGFT  RVLIWFISI  RVLIWLINI  RWVMWFGDG  SFLIWLLDF  SFLIWLLLC  SIIIWVVWI  SLLIWVISL  SLLMWMLRL  SLMIWLQTF  SLVIWICLV  SMVIWLLGF  SVVIWWIVC  TFLMWFIET  TIIIWLFFL  TLQIWLRQA  TLQIWVIWL  TVLMWPRKI  TVMMWPLAV  TVVMWVSAS  VILIWISVL  VLLIWLLTL  VLLMWLLVL  VVIIWLFLA  VVVMWIILA  VWLIWFTGS  VWVMWMRGG  WLLIWLLLG |

List of potential cross-reactive peptides identified in the X-scanning for NY_1 and NY_2 DARPin candidates. T2 cells were pulsed with each of the peptides listed in the table and CD8^+^ T-cell activation (TCA) was evaluated (percentage intracellular IFN-γ, normalized to TCA of NY-ESO1_157–165_(9V)-pulsed T2 cells). Both DARPin candidates were tested on the full set of peptides identified. In the absence of TCA, no percentage value is given in the table. Green font: TCA for naturally occurring NY-ESO1_157–165_ peptide (SLLMWITQC). Red font: TCA in the range of than NY-ESO1_157–165_ peptide. Orange font: low level of TCA.

**Supplementary Table 3. EC_50_ values for different TCEs**

| DARPin variant | | EC_50_ [nM]  NY_1xCD3 | EC_50_ [nM]  NY_2xCD3 |
| --- | --- | --- | --- |
| Linker variant  (length) | L (38 AA)  M (24 AA)  S (18 AA)  XS (11 AA)  XXS (6 AA) | 1.2  0.4  0.4  0.3  0.1 | 2.9  1.2  1.1  0.4  0.3 |
| CD3ε-binder variant | variant-1 (v1)  variant-2 (v2)  variant-3 (v3) | 7.1  0.5  0.005 | 19.9  0.7  0.02 |

The EC_50_ values were obtained from the analysis of CD8^+^ T cell activation curves for DARPin TCEs with varying linker lengths and CD3ε-binder variants. See **Figures 4A and 4C**.

**Supplementary Table 4. Cryo-EM data collection, refinement and validation of the NY_1/HLA-A*0201/NY-ESO1_157-165_(9V) complex**

| EMDB ID  PDB ID | EMD-50336  9FE1 |
| --- | --- |
| **Data collection and processing** |  |
| Magnification | 130k |
| Voltage (kV) | 300 |
| Electron exposure (e–/Å^2^) | 57.5 |
| Defocus range (μm) | -0.6 to -2.5 |
| Pixel size (Å) | 0.648 |
| Symmetry imposed | C1 |
| Initial particle images (no.) | 7,726,585 |
| Final particle images (no.) | 204,743 |
| Half-map based resolution (Å) at FSC_0.143_  3DFSC global resolution (Å) / sphericity | 3.0  3.1 / 0.74 |
| **Refinement** |  |
| Initial models used (PDB code) | 9EPA and 1S9W |
| Map sharpening *B_iso_* factor (Å^2^) | 143.4 |
| Model composition  Non-hydrogen atoms  Protein residues  Ligands  Waters | 8464  539  0  0 |
| *B* factors (Å^2^)  Protein (min / max / mean) | 23.55 / 112.13 / 43.78 |
| R.m.s. deviations  Bond lengths (Å)  Bond angles (°) | 0.005  0.849 |
| Validation  MolProbity score  Clashscore  Poor rotamers (%) | 1.82  9.57  0.22 |
| Ramachandran plot  Favored (%)  Allowed (%)  Disallowed (%) | 95.48  4.52  0.00 |
| **Map-model correlation and resolution estimates**  dFSC_model_ (Å)  d_mode_**_l_** (Å)  CC_mask_ (0.5-0.7 for low-high correlation)  CC_box_  CC_peaks_  CC_volume_ | sharpened / density-mod  3.5 / 2.9  3.2 / 2.9  0.70 / 0.68  0.77 / 0.58  0.67 / 0.58  0.72 / 0.67 |

**Supplementary Table 5. Statistics of the crystal structure of DARPin NY_1**

| PDB ID | 9EPA |
| --- | --- |
| **Data collection and processing** |  |
| Resolution range | 28.48 - 1.761 (1.824 - 1.761) |
| Space group | C 1 2 1 |
| Unit cell | 106.558 43.284 32.409 90 93.784 90 |
| Total reflections | 95069 (8028) |
| Unique reflections | 14698 (1438) |
| Multiplicity | 6.5 (5.6) |
| Completeness (%) | 99.55 (96.77) |
| Mean I/sigma(I) | 6.53 (2.17) |
| Wilson B-factor | 18.22 |
| R-merge | 0.2024 (0.6391) |
| R-meas | 0.2204 (0.7047) |
| R-pim | 0.08609 (0.2918) |
| CC1/2 | 0.982 (0.796) |
| **Refinement** |  |
| Reflections used in refinement | 14691 (1437) |
| Reflections used for R-free | 1460 (146) |
| R-work | 0.1835 (0.2392) |
| R-free | 0.2156 (0.2940) |
| CC(work) | 0.960 (0.889) |
| CC(free) | 0.971 (0.845) |
| Number of non-hydrogen atoms | 1276 |
| macromolecules | 1154 |
| ligands | 0 |
| solvent | 122 |
| Protein residues | 157 |
| RMS bonds/angles | 0.007 Å / 0.93° |
| Ramachandran favored/allowed (%) | 98.71/1.29 |
| Ramachandran outliers (%) | 0.00 |
| Rotamer outliers (%) | 0.00 |
| Clashscore | 3.40 |
| Average B-factor | 20.21 |
| macromolecules | 19.31 |
| solvent | 28.71 |

**Supplementary Table 6. Antibodies used for flow cytometry analyses**

| **Antibody** | **Cat# + company** |
| --- | --- |
| Live Dead stain aqua | L34957, Thermo |
| Live/Dead stain FITC | L23101, Thermo |
| APC mouse anti-human IFN-g | 554702, BD |
| Pacific Blue mouse anti-human CD8 | 558207, BD |
| CD8 Alexa488 | 557696, BD |
| CD25 PerCP Cy5.5 | 45-0259-42, ebio |
| CD69 Pe-Cy7 | 560712, BD |
| CD4 efluor 450 | 48-0048-42, ebio |
| CD3 PE | 12-0037-42, ebio |
| Penta-His AF488 | 35310, Qiagen |

**Supplementary Table 7. Tumor cell lines**

| **Cell lines** | **Source / number** |
| --- | --- |
| T2  MCF-7  U266B1  IM9  Colo-205  HCT116  NCI-H1755  NCI-H1703  MDA-MB231 | ATCC/CRL-1992  ATCC/HTB-22  ATCC/TIB-196  ATCC/CCL-159  ECACC 87061208  ATCC CCL-247  ATCC/CRL-5892  ATCC/CRL-5889  ATCC/HTB-26 |

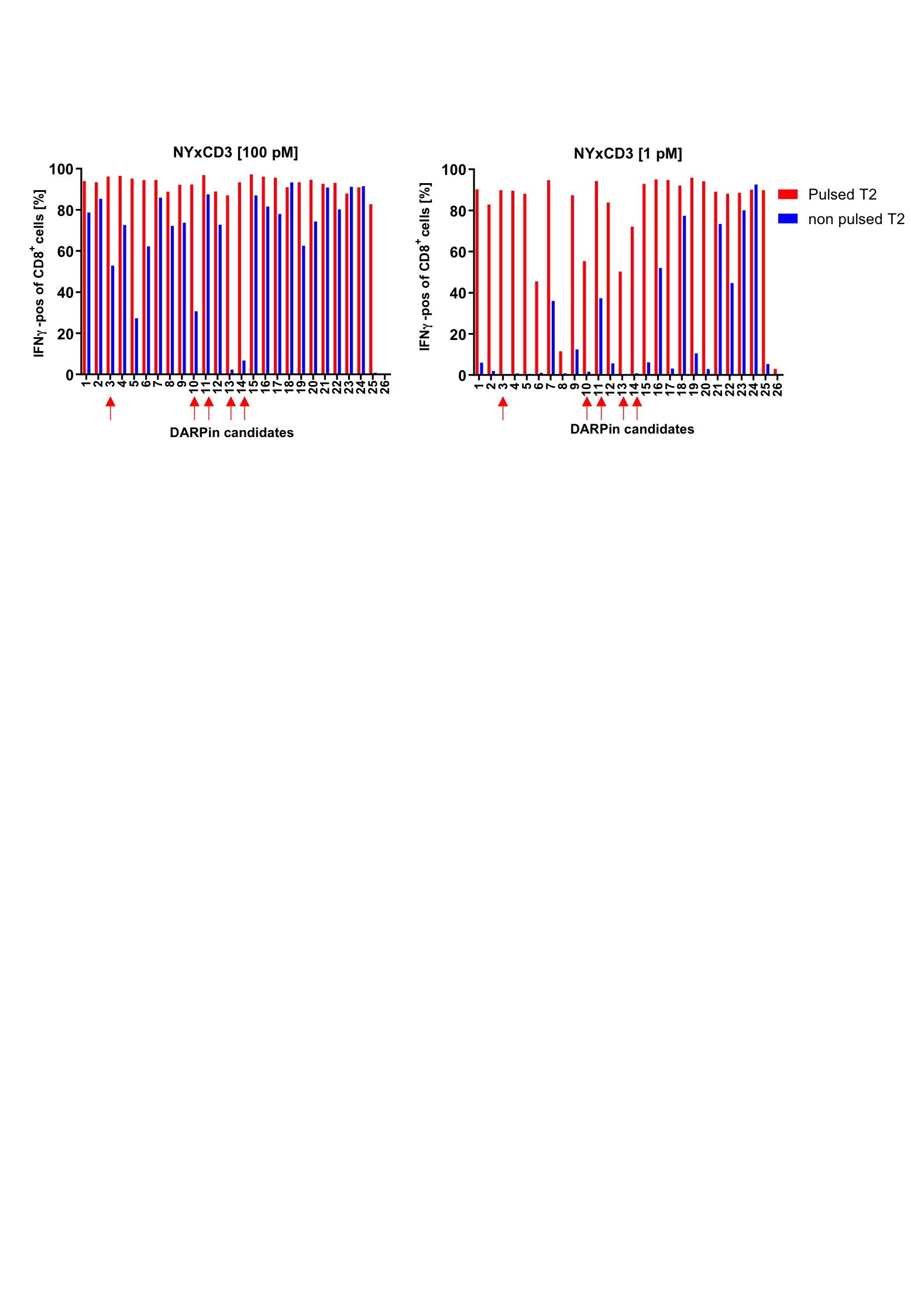
**Supplementary Figure 1. Functional screening of DARPin TCEs using NY-ESO1_157-165_(9V)-pulsed T2 cells**

26 different DARPin TCEs were tested for their capacity to elicit intracellular IFNγ release in CD8^+^ T cells in the presence of T2 cells pulsed with 1 and 100 pM NY-ESO1_157-165_(9V). The effects of peptide-pulsed T2 cells (red) on IFNγ production by CD8^+^ T cells were compared to the effects of non-pulsed (blue) T2 cells. DARPin TCEs that provoked significant IFNγ release by CD8^+^ T cells in the presence of peptide-pulsed T2 cells, and none in the presence of non-pulsed T2 cells were selected for further studies (indicated by red arrows).

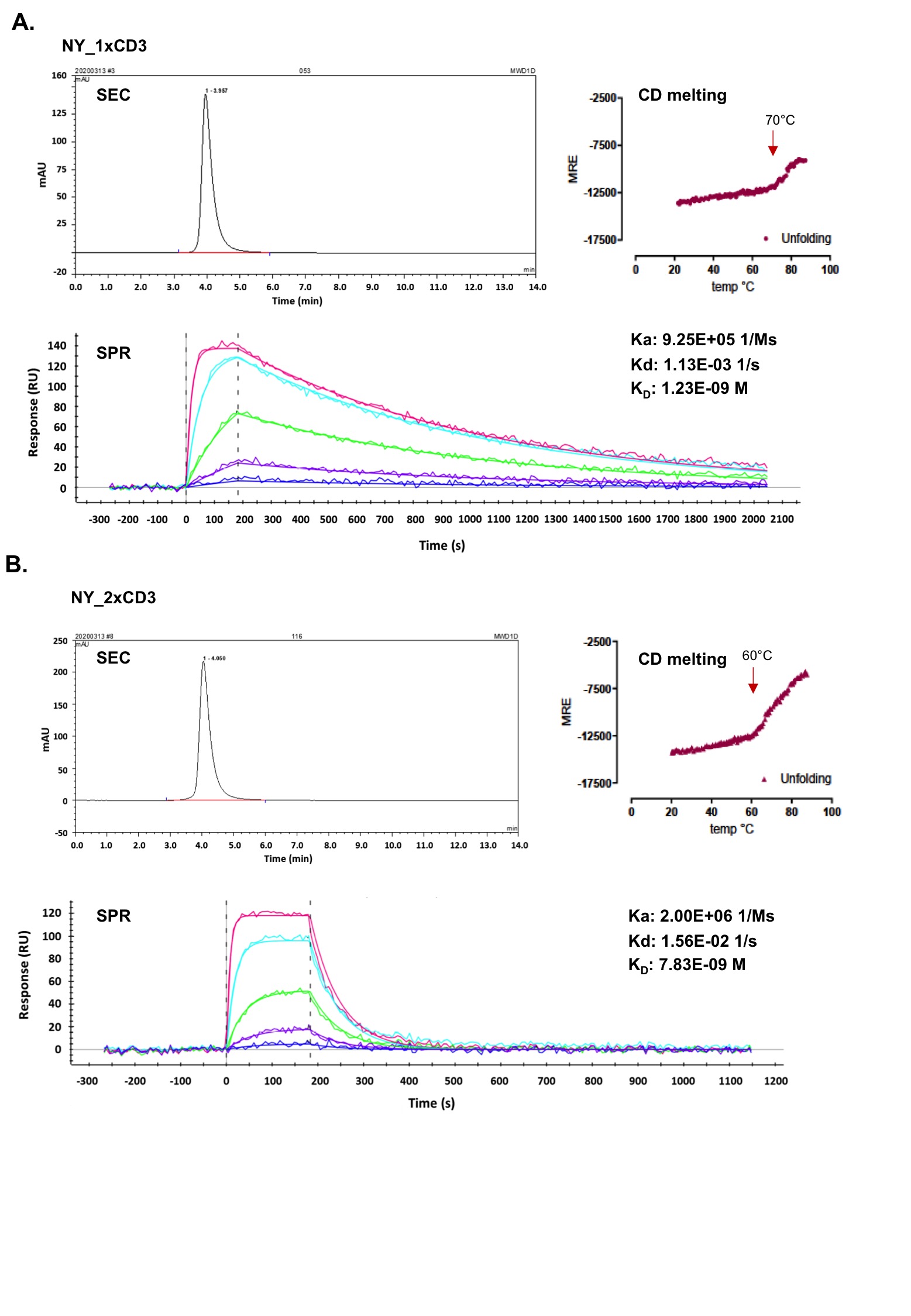
**Supplementary Figure 2. Biophysical characterization of the lead DARPin TCEs NY_1xCD3 and NY_2xCD3**

Biophysical analyses of **A.** NY_1xCD3 and **B.** NY_2xCD3 characterize both DARPin TCEs as homogenous, stable and with high affinity to HLA-A*0201/NY-ESO-1_157-165_(9V). Size exclusion chromatography (SEC) reveals single peaks, indicating homogeneous populations. Circular dichroism (CD) melting curves demonstrate that the two TCEs display high stability with T_m_ values at 70°C and 60°C for NY_1xCD3 and NY_2xCD3, respectively. Surface plasmon resonance (SPR) analyses show similar nanomolar affinities to HLA-A*0201^+^/NY-ESO1_157-165_(9V), but different kinetics.

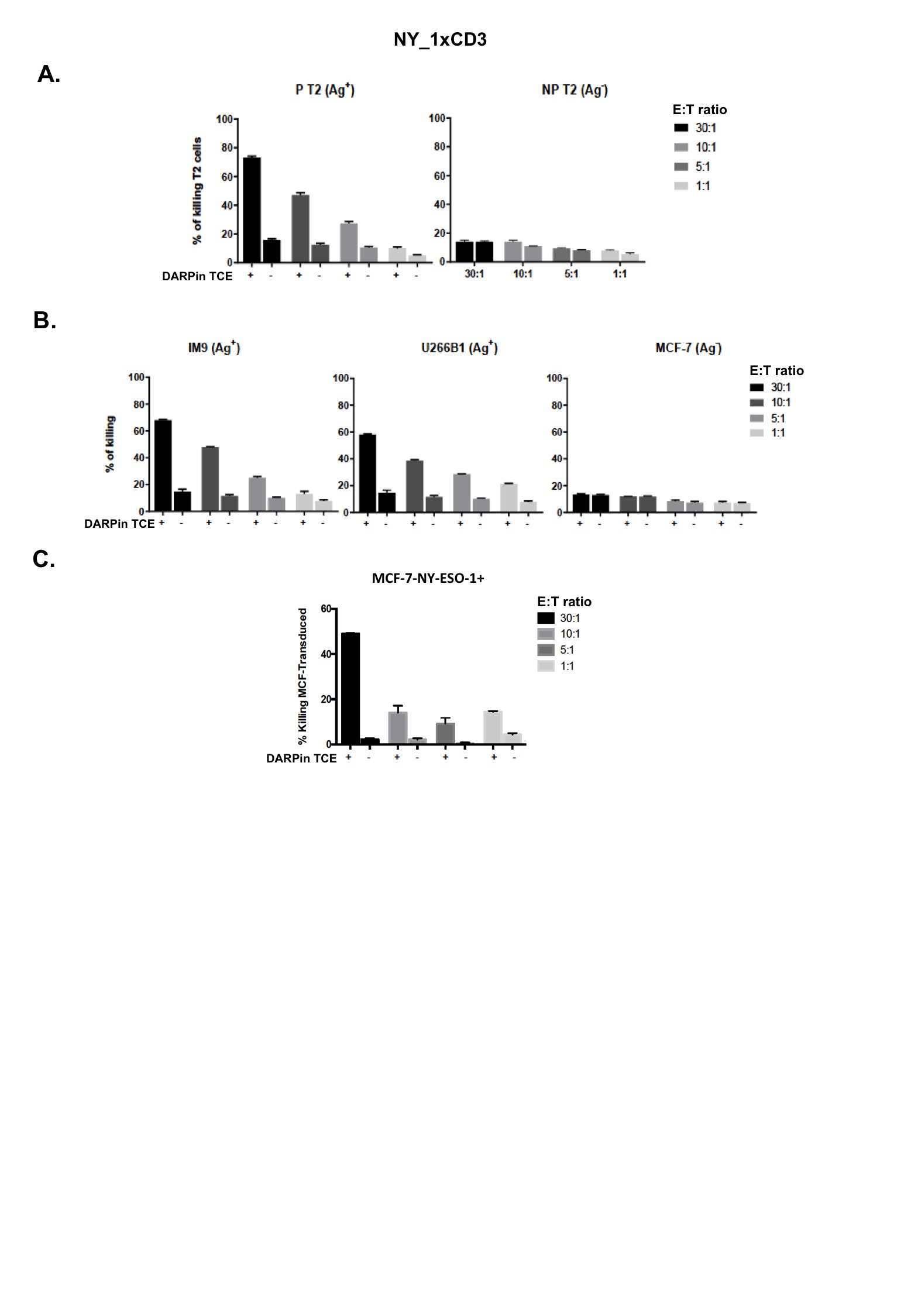
**Supplementary Figure 3. NY_1xCD3 mediates highly specific and efficient cytotoxicity towards HLA-A*0201^+^/NY-ESO1_157-165_^+^ tumor cell lines**

**A.** NY_1xCD3 enhances significantly the killing of T2 cells pulsed with 1 μM NY-ESO1_157-165_ by endogenous CD8^+^ T cells. The addition or not of NY_1xCD3 (10 nM) is indicated by + and -, respectively. E:T ratios stand for CD8 T cell effector:target cells. The percentage of specific lysis obtained for the different tumor cell lines by the chromium release assay are presented. **B.** NY_1xCD3 provokes significantly more efficient killing of HLA-A*0201^+^/NY-ESO1_157-165_^+^ but not HLA-A*0201^+^/NY-ESO1_157-165_^-^ cancer cell lines. **C.** NY_1xCD3 provoke efficient killing of the HLA-A*0201^+^/NY-ESO1_157-165_^-^ cancer cell line MCF-7 following transfection with full-length NY-ESO1 molecule.

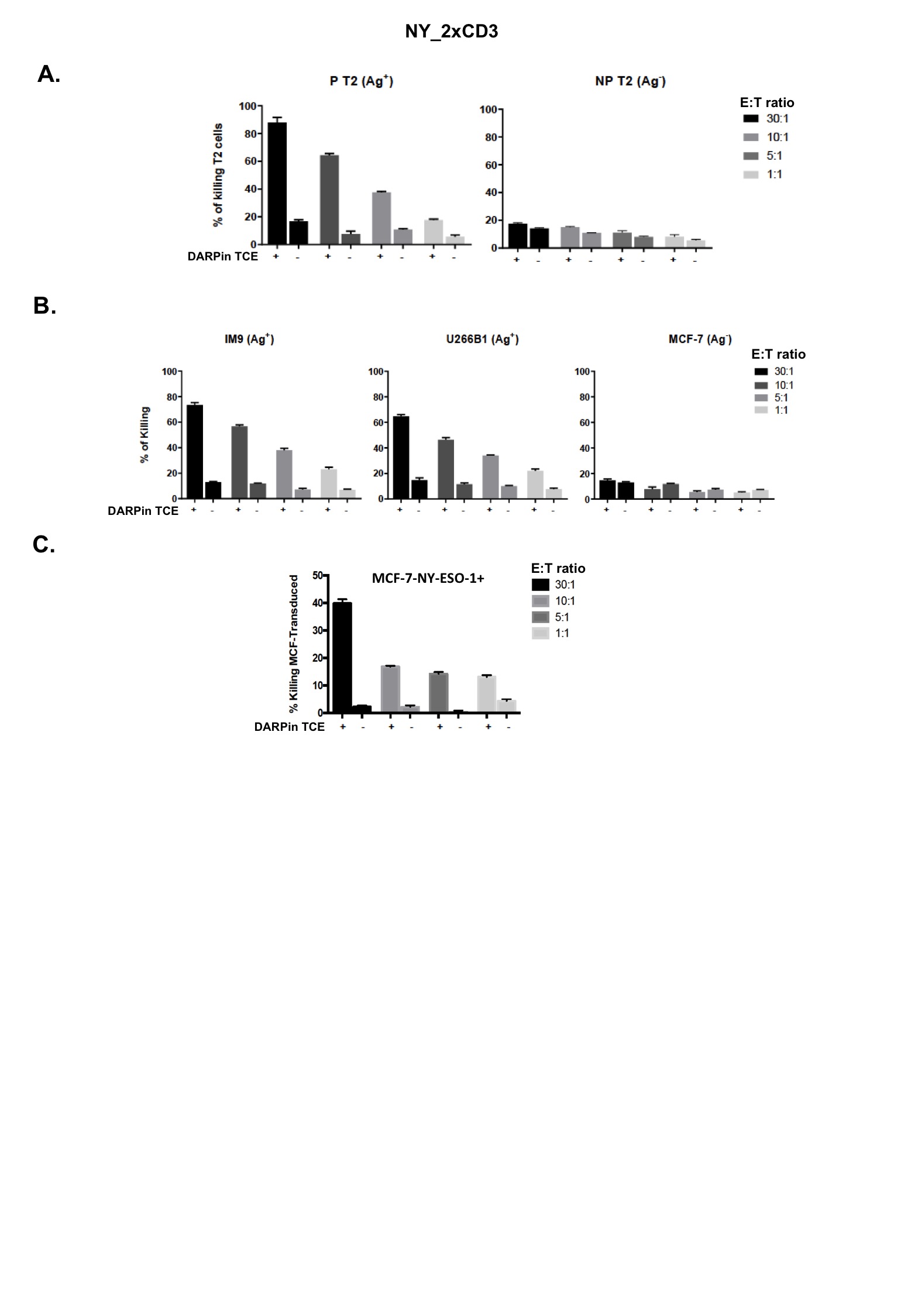
**Supplementary Figure 4. NY_2xCD3 mediates highly specific and efficient cytotoxicity towards HLA-A*0201^+^/NY-ESO1_157-165_^+^ tumor cell lines**

**A.** NY_2xCD3 enhances significantly the killing of T2 cells pulsed with 1 μM NY-ESO1_157-165_ by endogenous CD8^+^ T cells. The addition or not of NY_1xCD3 (10 nM) is indicated by + and -, respectively. E:T ratios stand for CD8 T cell effector:target cells. The percentage of specific lysis obtained for the different tumor cell lines by the chromium release assay are shown. **B.** NY_2xCD3 provokes significantly more efficient killing of HLA-A*0201^+^/NY-ESO1_157-165_^+^ but not HLA-A*0201^+^/NY-ESO1_157-165_^-^ cancer cell lines. **C.** NY_2xCD3 provoke efficient killing of the HLA-A*0201^+^/NY-ESO1_157-165_^-^ cancer cell line MCF-7 following transfection with full-length NY-ESO1 molecule.

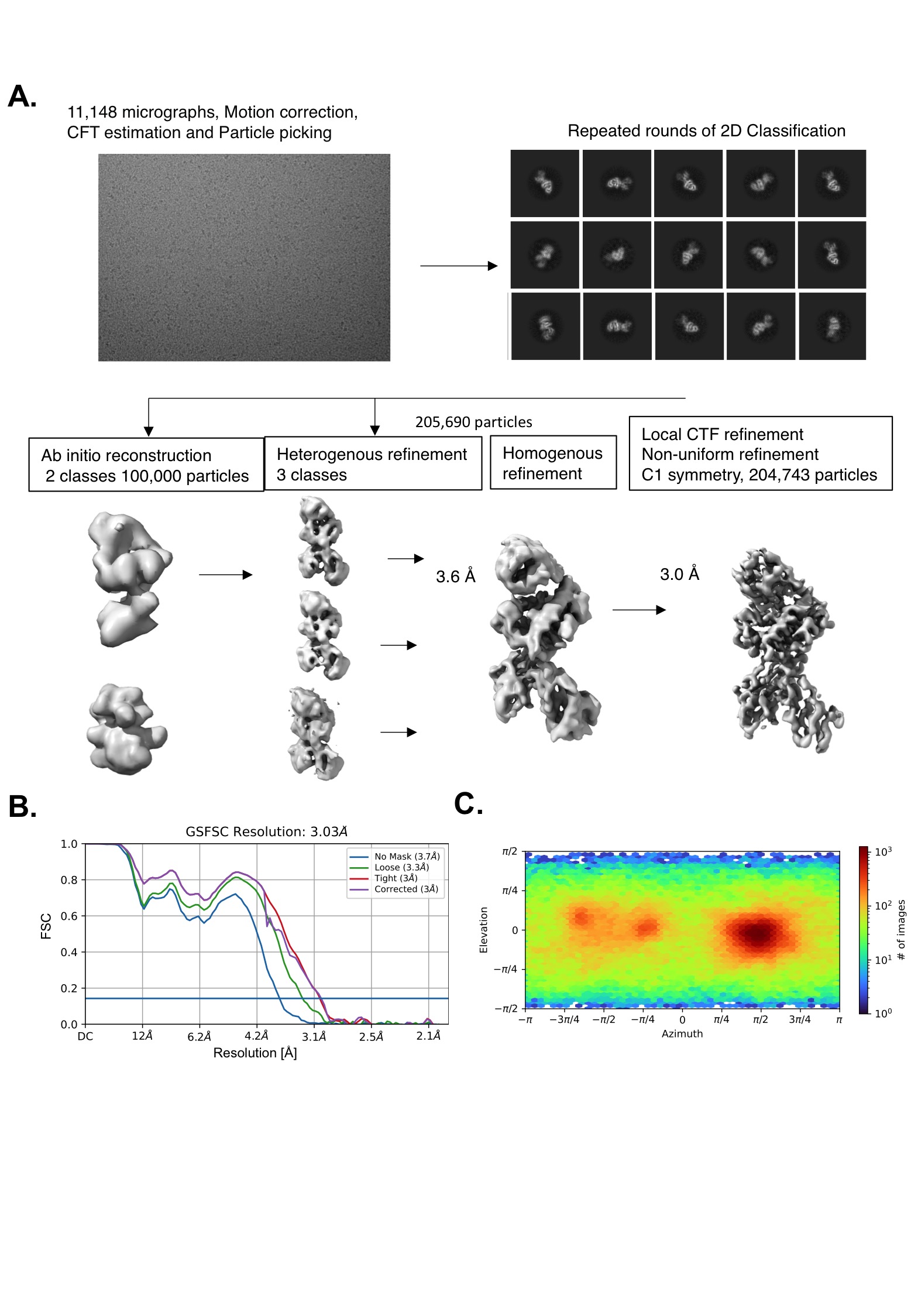
**Supplementary Figure 5. Cryo-EM processing scheme for the determination of the ternary NY_1/HLA-A*0201^+^/NY-ESO1_157-165_ structure**

**A.** Cryo-EM data processing workflow. **B.** Overall resolution estimation by Fourier Shell Correlation (FSC) with threshold 0.143. **C.** Viewing direction distribution plot (obtained from cryoSPARC v3.2.0 ^1^).

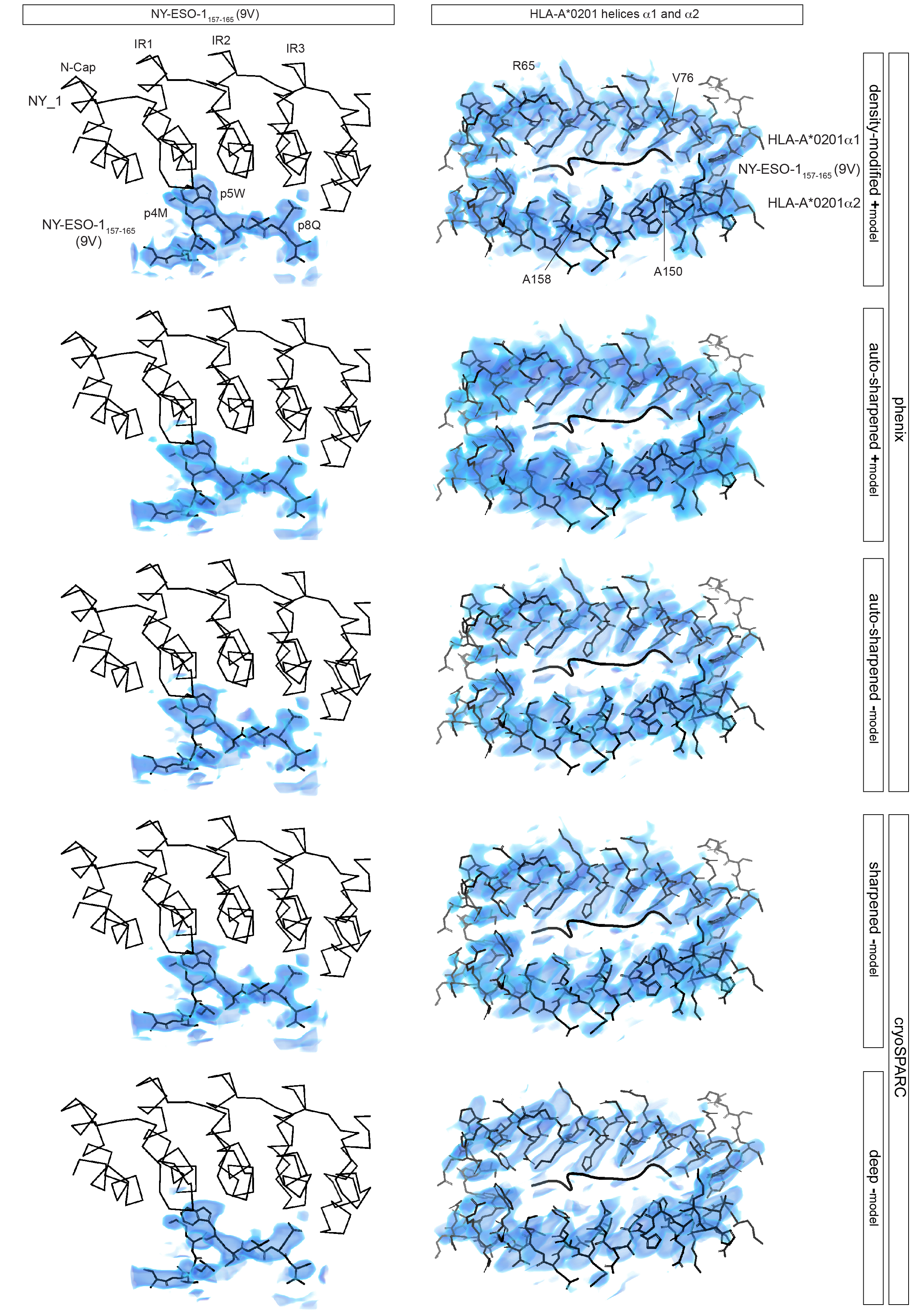
**Supplementary Figure 6. Map comparison of the regions defining the NY-ESO1_157-165_ peptide and the HLA-A*0201 α-helices**

Volumes of five post-processed maps are shown for the peptide NY-ESO1_157-165_(9V) and the two HLA-A*0201 α-helices in the same orientations as in **Figure 6D**, organized as row- and column-wise panels, respectively. From top to bottom, volumes were obtained after model-based density-modification (+), and auto-sharpening with (+) and without (-) model in Phenix as well as sharpening and deep-enhancement in cryoSPARC ^2-4^. Volume color ramp is identical to that in **Figure 6D**. Taking into account the additional map regions shown in **Supplementary Figures 6 and 7**, as well as the model-map FSC profiles, we selected the density-modified map as the final map for presentation.

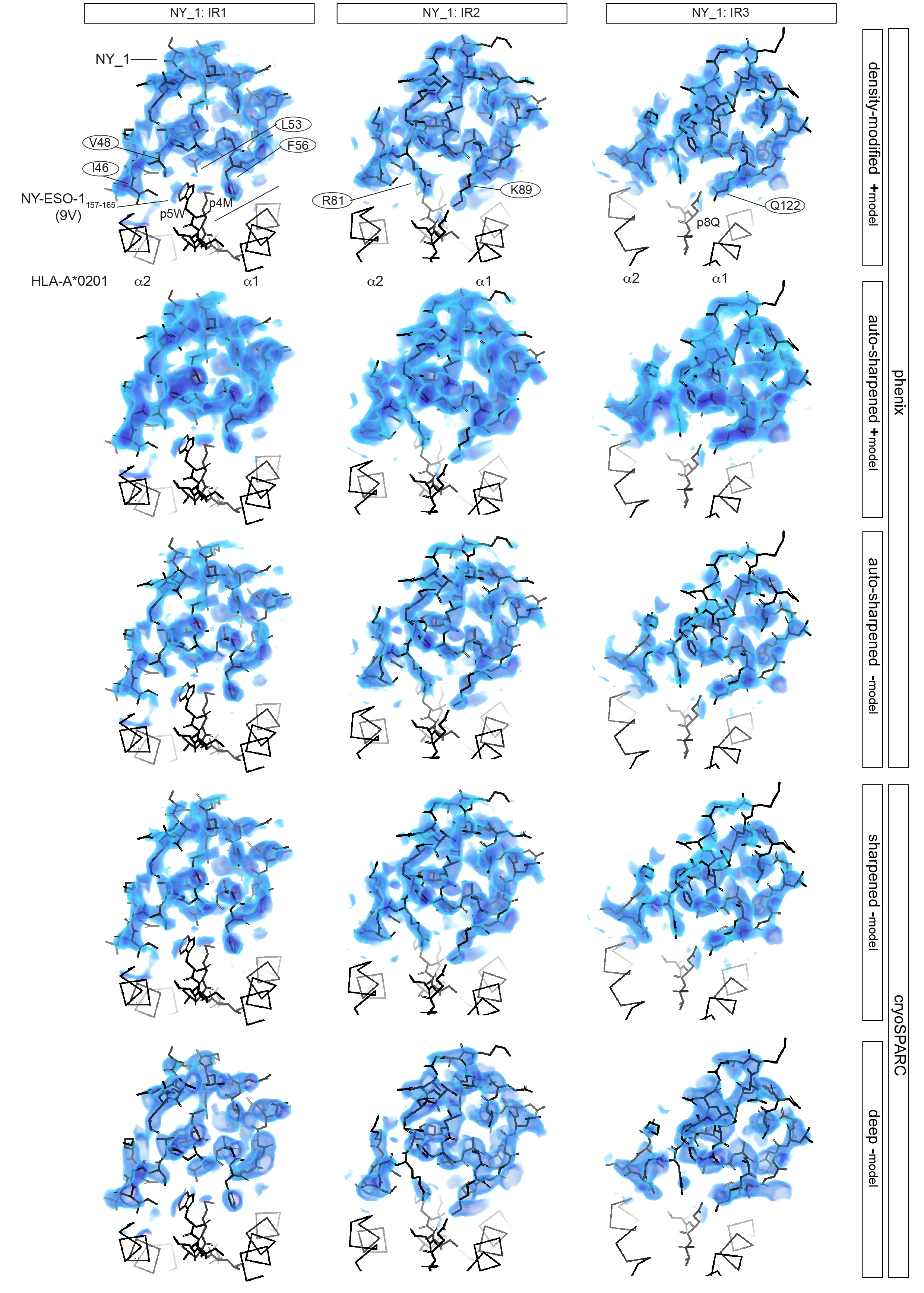
**Supplementary Figure 7. Map comparison of the regions defining internal repeats of NY_1**

Volumes of five post-processed maps are shown for the three internal repeats of NY_1 in the same orientations as in **Figure 6D**, organized as row and column-wise panels, respectively. Volumes obtained as described in **Supplementary Figure 5**.

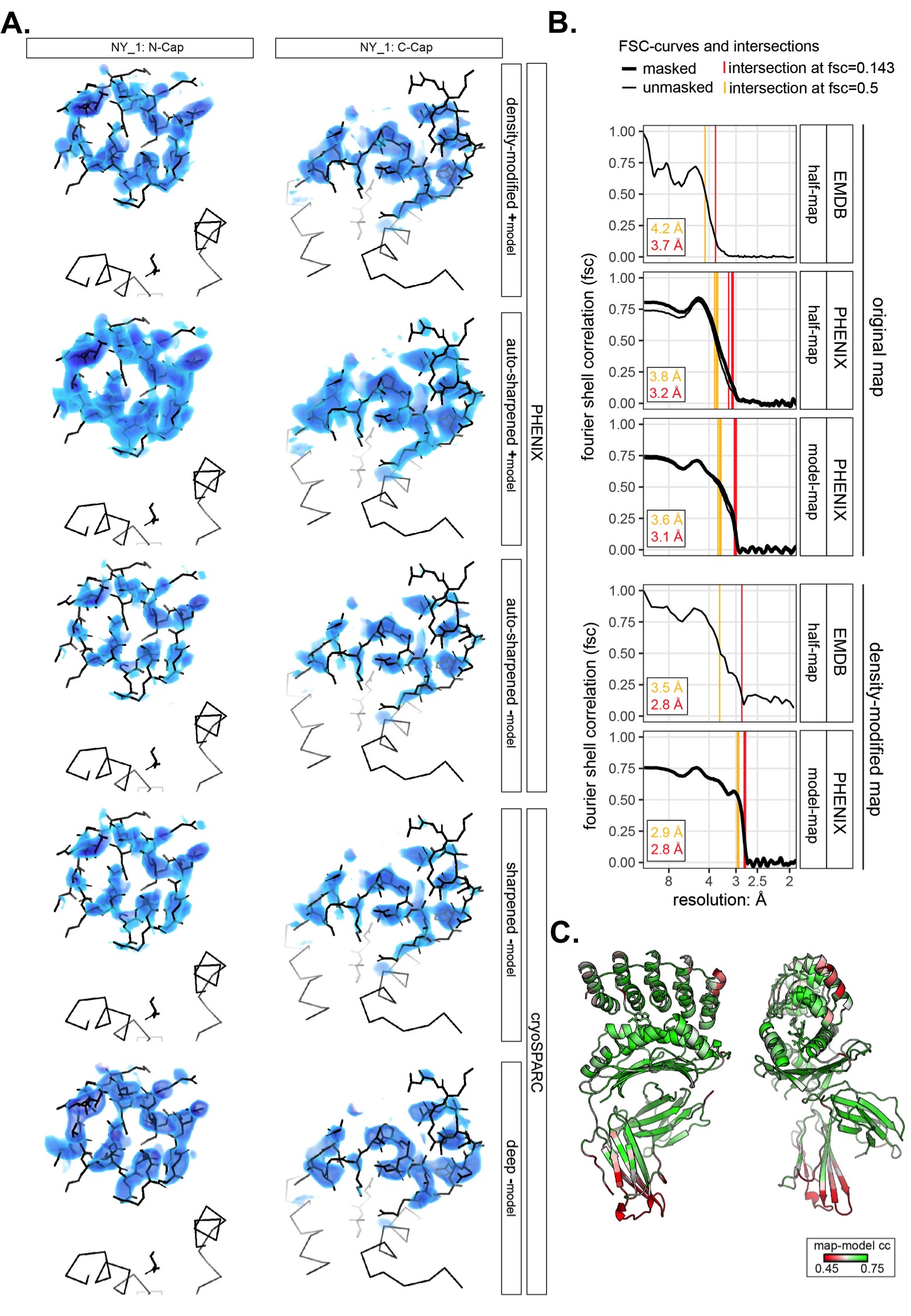

**Supplementary Figure 8. Map comparison of the regions defining the cap elements of NY_1, as well as presentation of half-map and model-map FSC curves and map-model cross-correlation**

**A.** Volumes of five post-processed maps are shown for the N- and C-Cap of NY_1 in the same orientations as in **Figure 6D**, organized as row and column-wise panels, respectively. Volumes were obtained as described in **Supplementary Figure 5**.

**B.** Half-map and model-map Fourier Shell Correlation (FSC) curves for the initial EMD-50336 deposited map and the five post-processed maps, respectively. The FSC curve of the masked half-maps crosses the value of 0.143 (red lines) at a resolution of 3.1 Å. For the five maps, the model-map FSC curves cross the value of 0.5 (orange lines) at resolutions of 2.9, 3.5, 3.5, 3.5 and 3.9 Å from top to bottom, respectively. The alternate model-based resolution estimate d_model_ ^5^ gives values of 2.9, 3.2, 3.4, 3.3 and 3.3 Å, respectively. Curves related to masked and unmasked maps are displayed as solid and dotted lines. **C.** The structure is colored in red, white and green for map-model cross-correlation values of 0.45, 0.6 and 0.75, respectively. Map-model cross-correlation values of 0.5 and 0.7 indicate poor and good fits ^5^. The views are identical to those in **Figures 6A and 6B**.

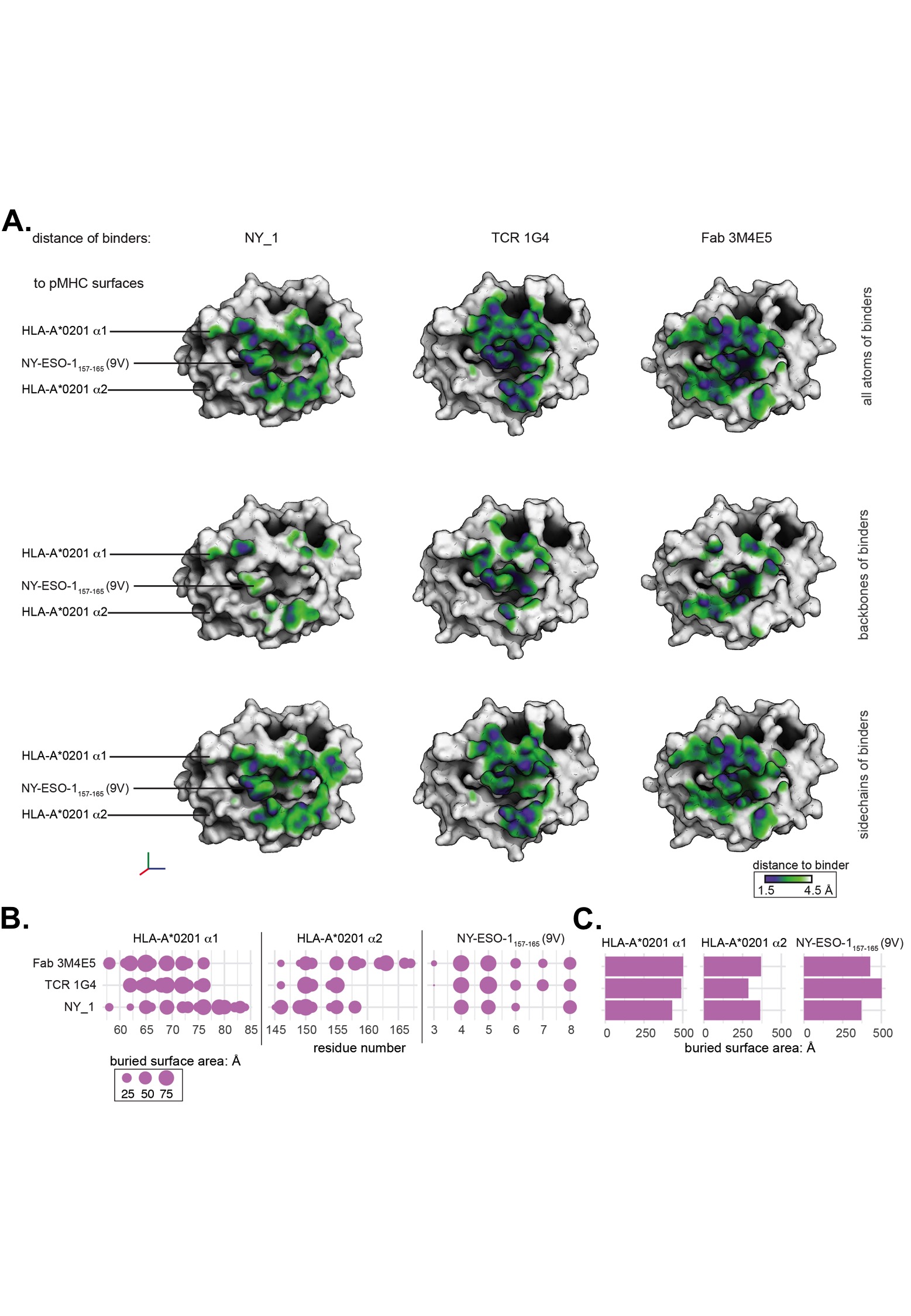
**Supplementary Figure 9. In contrast to NY_1, the TCR 1G4 and the Fab fragment 3M4E5 bind to HLA-A*0201/NYESO1_157-165_(9V) with tighter interfaces and closer binder-to-target distances**

**A.** The distance ramps between DARpin NY_1 (left panel), TCR 1G4 (middle) and Fab fragment 3M4E5 (right), and HLA-A*0201/NY-ESO1_157-165_(9V) are presented for all atoms (top panel), backbone atoms (middle panel) and side chains (bottom panel). This structural comparison demonstrates that the CDRs of both 1G4 and 3M4E5 form tighter interfaces with the surface of HLA-A*0201/NY-ESO1_157-165_(9V), compared to the binding mode and interface used by NY_1.

**B.** Residue-level buried surface areas were obtained from PISA ^6^ and plotted separately for each binder in panels for the HLA-A*0201 helices α1 and α2 as well as the NY-ESO1_157-165_(9V) peptide. The areas of the filled circles scale with the buried surface area (Å^2^).

**C.** The residue-level buried surface areas are summed for each binder and the panels of HLA-A*0201 helices α1 and α2 as well as NY-ESO1_157-165_(9V).

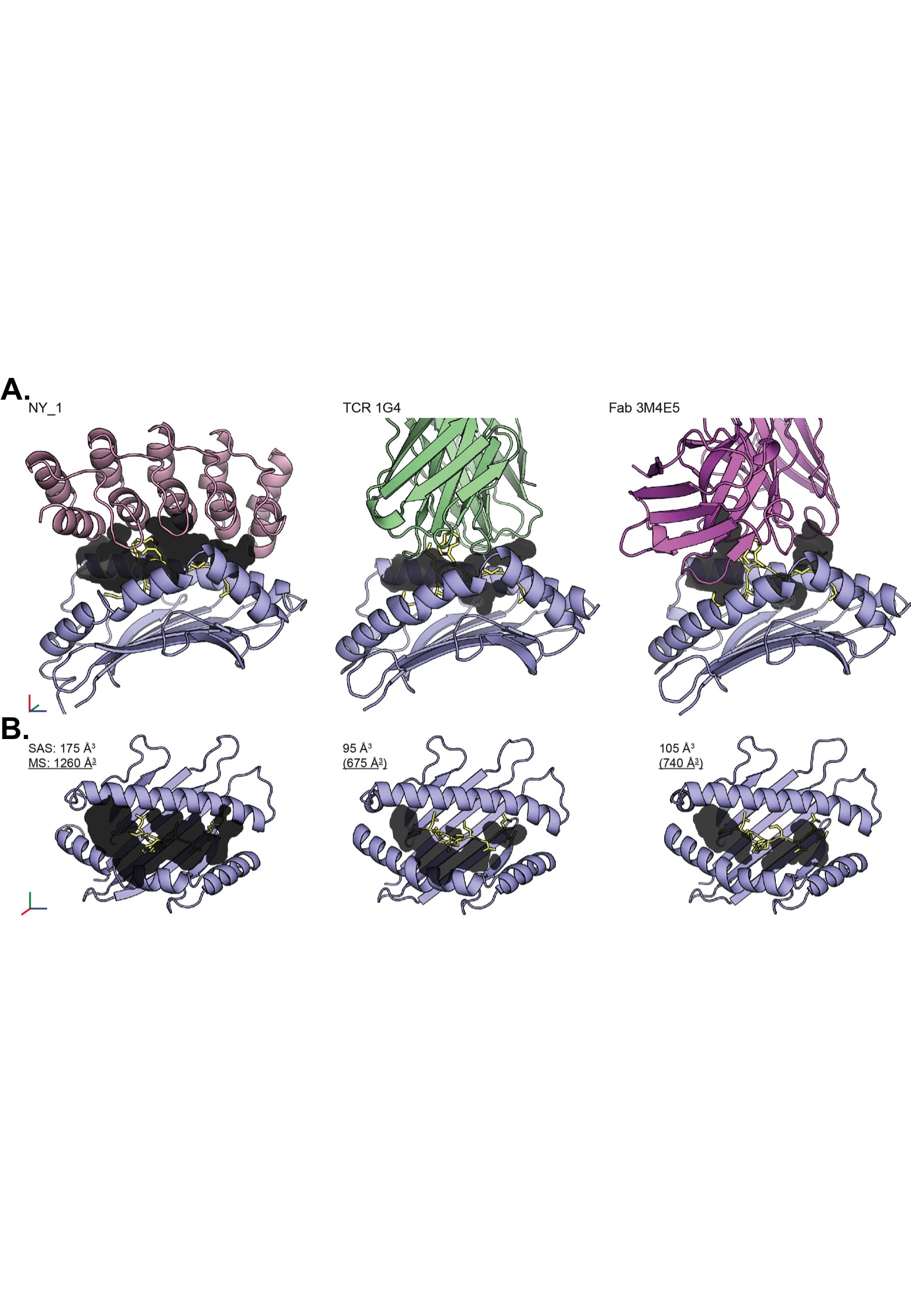

**Supplementary Figure 10. The NY_1/HLA-A0201/NYESO1_157-165_(9V) interface comprises a significantly larger void compared to those formed with TCR 1G4 or Fab 3M4E5**

**A.** Voids in contact with the peptide for complexes of HLA-A0201/NY-ESO1_157-165_(9V) with DARPin NY_1, TCR 1G4 and Fab fragment 3M4E5 were obtained using CastP ^7^. **B.** The summed volumes based on solvent-accessible surfaces (SAS) molecular surfaces (MS) are listed besides each complex. The analyses were performed only for surfaces created by atoms localized within 8Å around the peptide, and applying a default probe radius of 1.4Å. The voids are shown in dark within both the side- and top-views.

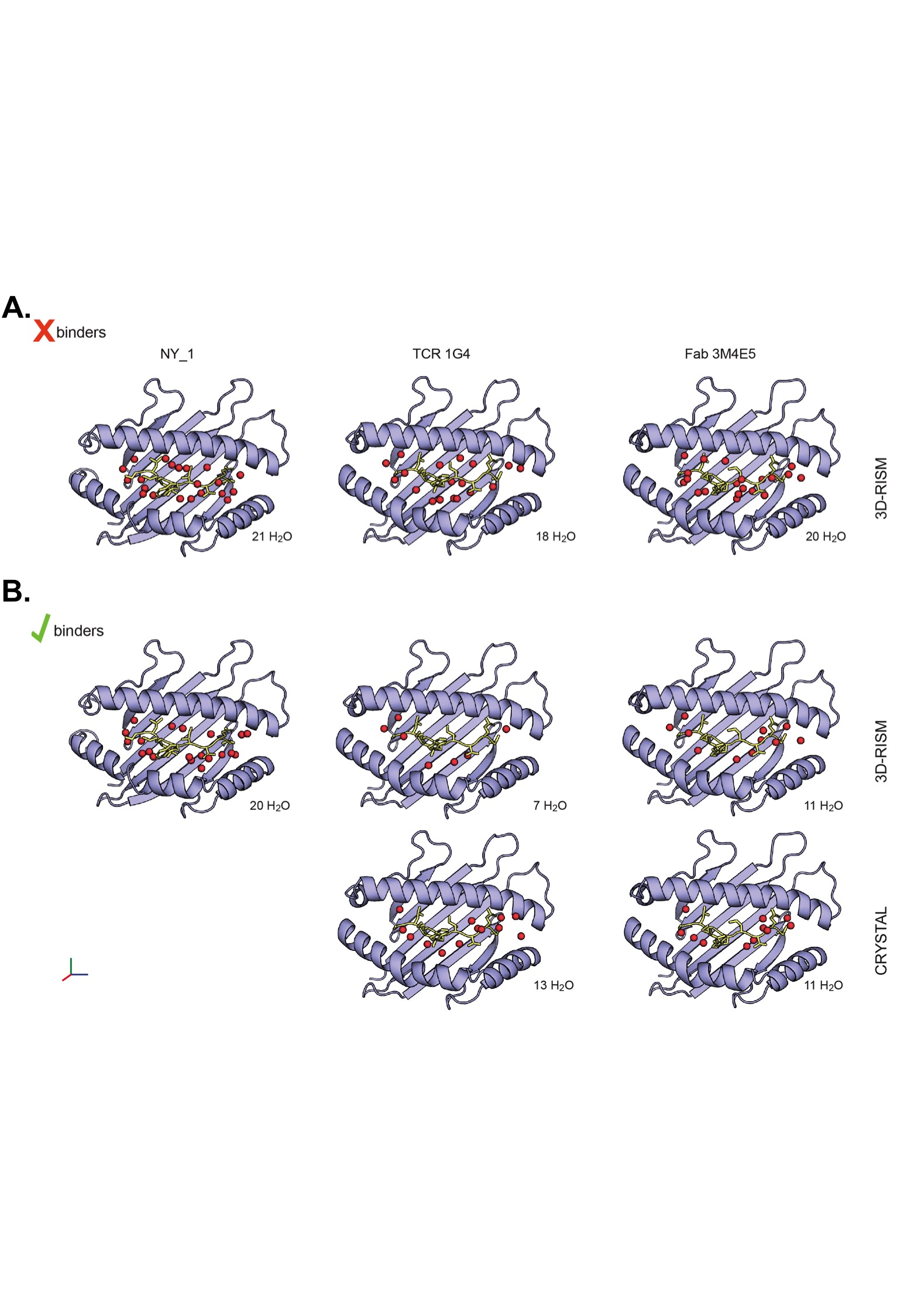

**Supplementary Figure 11. A larger amount of water molecules is predicted to remain within the NY_1/HLA-A*0201/NY-ESO1_157-165_(9V) interface after binding to the DARPin molecule**

**A.** Predictions using 3D-RISM ^8^ were made for HLA-A*0201/NY-ESO1_157-165_(9V) prior to binding to NY_1, 1G4 and 3M4E5, revealing similar amounts of H_2_O molecules surrounding the presented peptide. **B.** 3D-RISM predictions of the position and amount of water molecules at the interfaces formed between HLA-A*0201/NY-ESO1_157-165_(9V) and NY_1 (left), 1G4 (middle) and 3M4E5 (right) indicated that water molecules remain within the NY_1/HLA-A*0201/NY-ESO1_157-165_(9V). Water molecules found within the interfaces of the 1G4/HLA-A*0201/NY-ESO1_157-165_(9V) and 3M4E5/HLA-A*0201/NY-ESO1_157-165_(9V) interfaces are displayed for comparison.

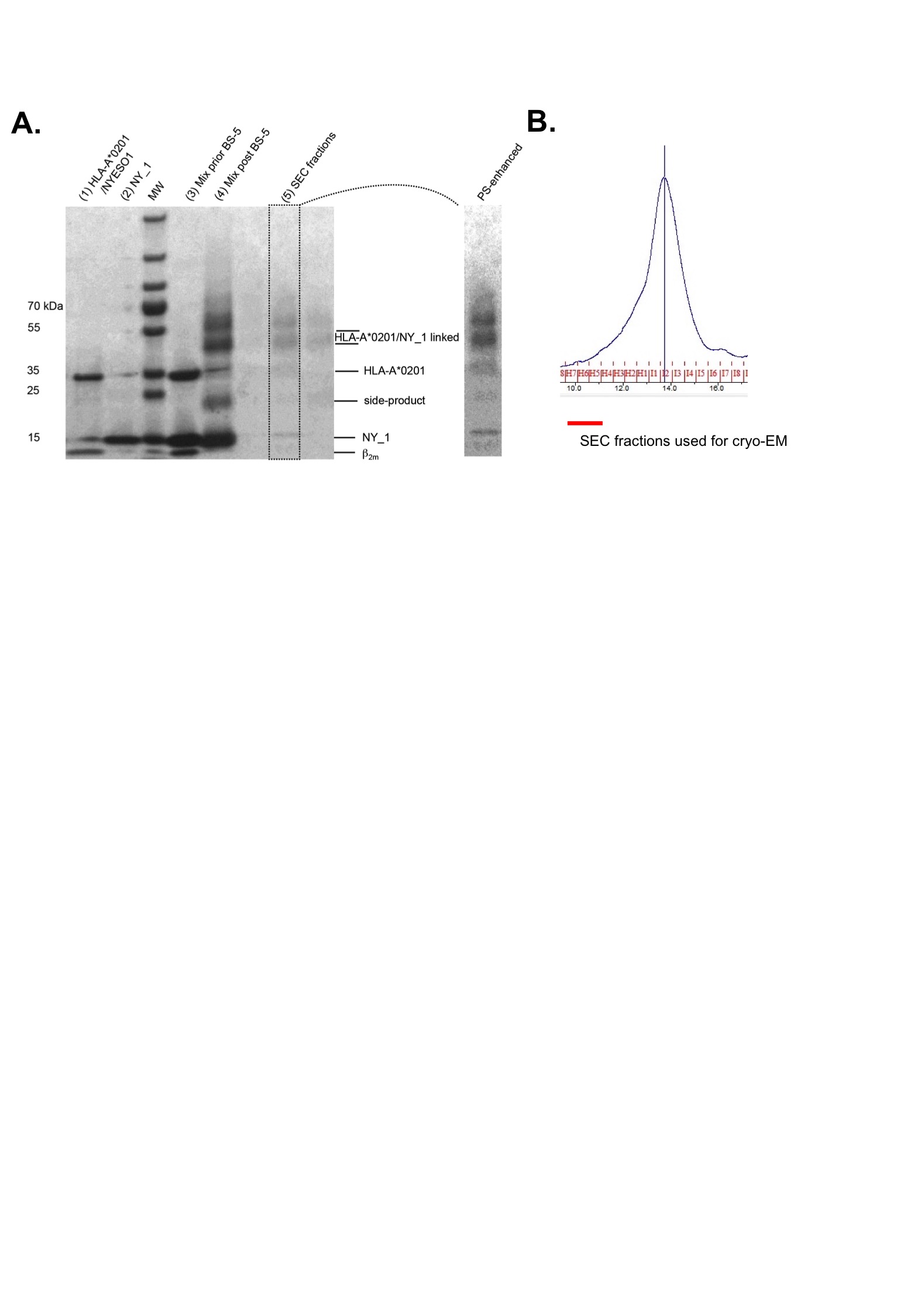
**Supplementary Figure 12. Production and isolation of cross-linked DARPin NY_1/HLA-A*0201/NY-ESO1_157-165_(9V) complexes**

**A.** Sample preparation for collection of the high-resolution single particle cryo-EM dataset. The following samples were applied: (1) SEC and Streptactin-purified refolded HLA-A*0201/NY-ESO1_157-165_(9V) (2) NiNTA- and SEC-purified NY_1 (3) 1:2 molar mix of HLA-A*0201/NY-ESO1_157-165_(9V) and NY_1 prior (4) and post BS5 cross-linking, as well as the (5) final combined SEC-fractions used for cryo-EM. To make the bands in the fifth lane more visible, the selected region was enhanced in Adobe Photoshop. Denaturing SDS-PAGE gradient gels were stained using silver blue ^9^. **B.** The cross-linked sample was purified by Superdex 200 10/300 to isolate the HLA-A*0201/NY-ESO1_157-165_(9V)/NY_1 complex. The fraction corresponding to the main peak eluted after 14 mL and was concentrated for grid preparation.
